# Integrated multiomic analysis identifies *TRIP13* as a mediator of alveolar epithelial type II cell dysfunction in idiopathic pulmonary fibrosis

**DOI:** 10.1101/2024.04.05.588292

**Authors:** Laurence St. Pierre, Asres Berhan, Eun K. Sung, Juan R. Alvarez, Hongjun Wang, Yanbin Ji, Yixin Liu, Haoze Yu, Angela Meier, Kamyar Afshar, Eugene M. Golts, Grace Y. Lin, Alessandra Castaldi, Ben A. Calvert, Amy Ryan, Beiyun Zhou, Ite A. Offringa, Crystal N. Marconett, Zea Borok

## Abstract

Idiopathic pulmonary fibrosis (IPF) is a lethal progressive lung disease urgently needing new therapies. Current treatments only delay disease progression, leaving lung transplant as the sole remaining option. Recent studies support a model whereby IPF arises because alveolar epithelial type II (AT2) cells, which normally mediate distal lung regeneration, acquire airway and/or mesenchymal characteristics, preventing proper repair. Mechanisms driving this abnormal differentiation remain unclear. We performed integrated transcriptomic and epigenomic analysis of purified AT2 cells which revealed genome-wide alterations in IPF lungs. The most prominent epigenetic alteration was activation of an enhancer in thyroid receptor interactor 13 (*TRIP13*), coinciding with *TRIP13* upregulation. *TRIP13* is broadly implicated in epithelial-mesenchymal plasticity and transforming growth factor-β signaling. In cultured human AT2 cells and lung slices, small molecule TRIP inhibitor DCZ0415 prevented acquisition of the mesenchymal gene signature characteristic of IPF, suggesting TRIP13 inhibition as a potential therapeutic approach to fibrotic disease.

## INTRODUCTION

Idiopathic pulmonary fibrosis (IPF) is a distal lung disease characterized by fibrotic remodeling of the alveolar niche leading to progressive loss of lung function^(1)^. Current therapies only delay progression^(2)^, leaving lung transplant as the sole remaining option^(3, 4)^. Risk factors include older age, male sex, smoking history^(1)^, other inhalant exposures^(5)^ and in familial cases, inherited mutations^(6, 7)^ primarily in genes linked to alveolar epithelial type II (AT2) cell biology^(8–12)^, implicating AT2 cells in IPF pathogenesis^(13)^. Furthermore, mutations affecting telomere maintenance have been implicated in both familial and sporadic IPF^(14)^. Telomere shortening has been suggested as a triggering event for dysregulated transforming growth factor-β (TGF-β) signaling^(15)^. Indeed, ample evidence supports a role for TGF-β in promoting epithelial-to-mesenchymal plasticity (EMP) that is implicated in the pathogenesis of IPF^(16–18)^.

Single-cell RNA-sequencing (scRNA-seq) analyses of IPF lungs have identified abnormal epithelial cell types co-expressing alveolar and airway (e.g., basal) cell genes, and atypical transitional cells expressing mesenchymal markers^(19–22)^. Expression of conducting airway markers in human alveolar epithelial cells (AECs) can be replicated in organoid cultures and is stimulated by TGF-β^(21)^. However, the molecular pathways underlying such TGF-β-mediated abnormal cell fate transitions remain poorly defined.

All cells in the body arise from a single genome; cell fate and identity are accordingly encoded by cell type-specific *epi*genomes. Environmental factors such as tobacco smoke and pollutants, and intrinsic factors such as aging, all implicated in IPF pathogenesis, can alter the epigenome^(23, 24)^. The contribution of epigenetic alterations to IPF has been studied in the limited context of genes of high interest, particularly the role of DNA methylation at certain target genes^(7, 25, 26)^, and global imbalances in histone deacetylase activity^(27)^. To our knowledge, no detailed investigation of epigenomic dysregulation within resident alveolar epithelial populations in IPF has been performed. Epigenomic regulation in AT2 cells has largely been studied in the non-diseased AT2 cell population ^(28, 29,30)^. Observations that AT2 cells can undergo metaplastic shifts in fate identity during fibrosis^(19–21, 31)^ implicate AT2 cell epigenomic remodeling in disease development. Here we carried out an integrated transcriptomic and epigenomic analysis of AT2 cells from IPF and non-diseased lung to investigate the mechanisms underlying IPF-associated epithelial dysfunction.

## RESULTS

### IPF AT2 cells display genome-wide dysregulation in gene expression patterns

We performed bulk RNA-sequencing (RNA-seq) and assay for transposase-accessible chromatin-sequencing (ATAC-seq) on purified distal lung epithelial populations enriched for AT2 cells. Cells were obtained from lungs of 3 IPF patients undergoing transplantation and 3 donors with no prior history of chronic airway disease (hereafter referred to as “controls”). Donor characteristics including AT2 cell purities are listed in Table S1. RNA for five of the six samples met quality control requirements needed for successful RNA sequencing (Figure S1). We thus used 5 samples for transcriptomic analysis. All six samples met ATAC quality control requirements (Figure S2).

We performed differential transcriptome analysis using DeSeq2^(32)^ (Figure 1A). 1559 genes were significantly activated (false discovery rate (FDR)-corrected p-value < 0.05), whereas 1232 genes were significantly downregulated and/or silenced in AT2-IPF samples relative to AT2-controls (Figure 1B). Differentially expressed genes (Table S2) were overrepresented in pathways known to be elevated in IPF (Figure 1C)^(33–36)^. Relative gene expression levels for a limited set of genes as markers for AT2-control cells, epithelial to mesenchymal transition (EMT), and basal cells (given the presence of basal-like epithelial cells in IPF^(19, 21, 22)^) are highlighted in Figure 1D. AT2 cell marker genes were reduced, while EMT markers were elevated in AT2-IPF cells^(37, 38)^. Basal- and basaloid-associated cell markers were also elevated in AT2-IPF cells^(22)^.

**Figure 1:**
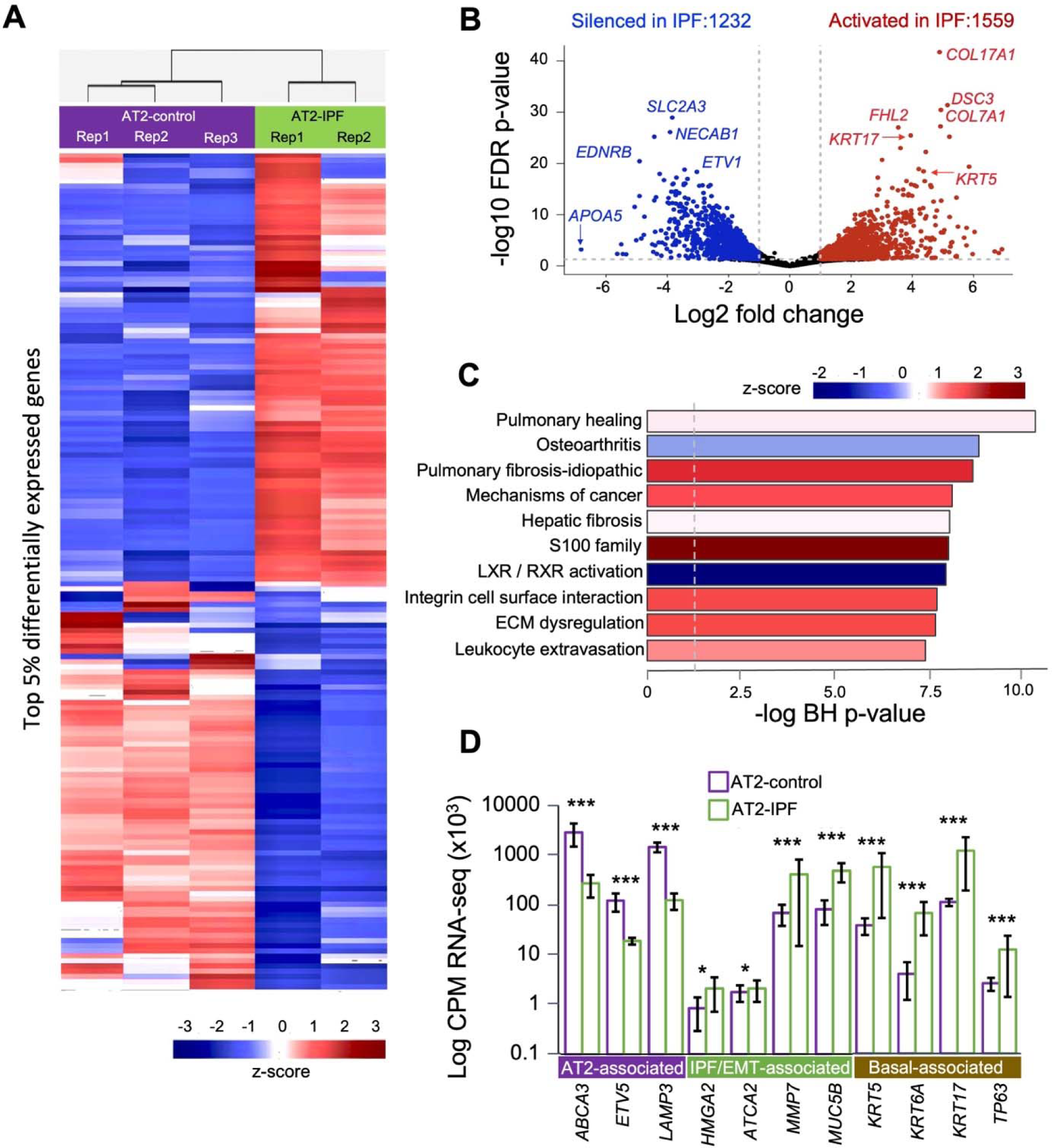
AT2-IPF cells exhibit genome-wide dysregulation in gene expression patterns. **A.** Supervised heatmap of top 5% significantly differentially expressed genes (false discovery rate (FDR) < 0.05) between AT2-control (purple) and AT2-IPF (green) samples. Expression counts normalized by z-score; red = elevated in IPF, blue = decreased in IPF. **B.** Volcano plot of differentially expressed genes between AT2-control and AT2-IPF. X-axis = log2-fold change between AT2-control and AT2-IPF. Y-axis = -log10 of the FDR-corrected p-value significance of gene expression changes. Red = activated in AT2-IPF relative to AT2-control, blue = decreased in AT2-IPF relative to AT2-control. Black = non-significant changes. Gray dotted lines = significance cutoffs. Select significant changes are indicated. **C.** Bar plot of pathway enrichment for differentially expressed genes in AT2-IPF vs. AT2-control generated in Ingenuity Pathways Analysis (IPA). X-axis = -log10 Benjamini-Hochberg corrected p-value of pathway enrichment significance. Bar color reflects Z-score for pathway enrichment; red = predicted activation in AT2-IPF, blue = predicted inhibition in AT2-IPF. **D**. Histogram of relative expression values for AT2-associated, EMT-associated and basal-associated genes in AT2-control (purple) vs. AT2-IPF cells (green).

### AT2-IPF cells display genome-wide epigenetic alterations

We performed bulk ATAC-seq on 5,000 to 50,000 cells per donor sample. ATAC libraries underwent next generation sequencing and realignment to the Hg38 genome. We first examined epigenomic variation between individual patients within AT2-control or AT2-IPF groups by assessing peak overlap for each group. AT2-control replicates shared over 16,000 accessibility peaks among all three samples, while AT2-IPF replicates shared over 24,000 peaks (Figure 2A). Replicates 1 of both the AT2-control and AT2-IPF samples were from current smokers (Table S1), perhaps accounting for some of the observed inter-sample variability.

**Figure 2:**
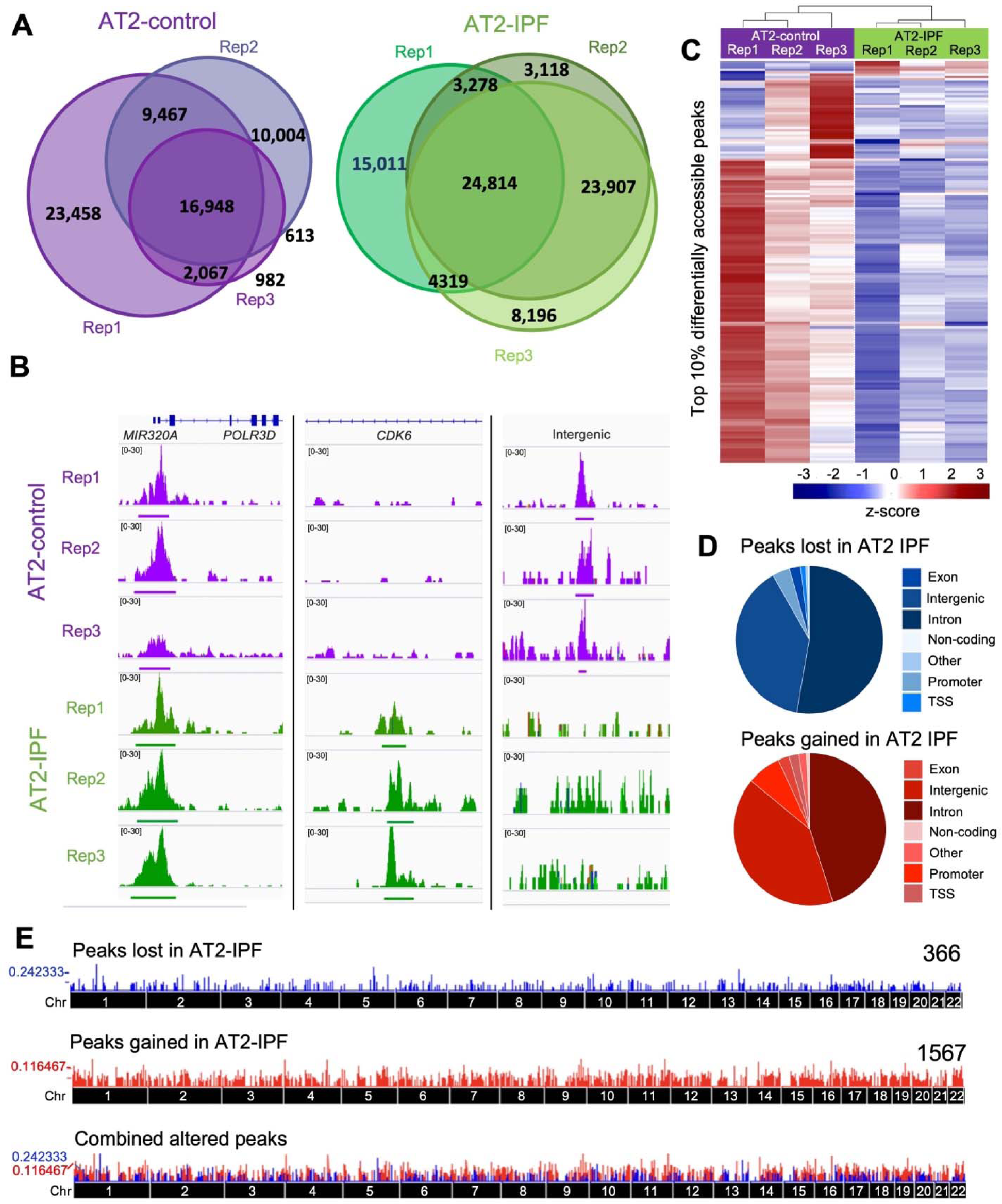
AT2-IPF cells exhibit genome-wide epigenetic alterations. **A.** Venn Diagrams of bulk ATAC-seq peak overlap among AT2-control (purple) and AT2-IPF (green) replicates. **B.** Peak enrichment for select loci displayed using Integrated Genomics Viewer (IGV). X = genomic locus peak occupies, y = peak height. AT2-control replicates = purple, AT2-IPF replicates = green. **C.** Heatmap of top 10% differentially accessible (DA) peaks, calculated as area under the curve (AUC) in GenRich. Ward clustering was used for samples (columns) and ATAC peak regions (rows). Red = enriched for ATAC reads within the peak, blue = little to no reads within the peak region. AT2-control = purple, AT2-IPF = green. **D.** Pie charts showing genomic distribution of consensus peaks with differential occupancy in AT2-control and AT2-IPF and their position relative to nearest-neighbor gene bodies. Regions closing (blue) and opening in IPF (red) are classified based on the indicated nearest neighbor gene structures: Exon = peak overlap with nearest neighbor-gene known exon region, Intergenic = peak occurs > 2.5 kb from transcriptional start site and outside of known gene regions, Intron = peak lies within known intron of nearest-neighbor gene, Non-coding = peak lies in the non-coding region of nearest-neighbor gene, Promoter = peak occurs within 2.5 kb of the transcription start site (TSS) of nearest-neighbor gene, TSS = peak lies within 1 kb of the TSS of nearest-neighbor gene, Other = peak overlaps with other known genomic elements (e.g., LINE, SINE, etc). Pie chart values are percentages of total peak occurrences. **E.** Genome graph plot of genomic distribution for peaks present in AT2-control and lost in AT2-IPF (blue, 366 peaks) and peaks gained in AT2-IPF (red, 1567 peaks). X-axis = human chromosomes 1-22 displayed as one contiguous line. Y-axis = number of peaks within a given region (∼250 k bases).

Examples of shared and differentially gained or lost peaks are shown (Figure 2B). A consensus set of peaks present in all three replicates was generated for the AT2-control and AT2-IPF samples. A heatmap of the top 10% differentially accessible peaks across the dataset shows marked changes in genome accessibility between the two groups, with the majority of top differential peaks consisting of those lost in AT2-IPF (Figure 2C). Each differentially accessible peak was next annotated to the nearest neighbor gene using HOMER^(39)^. The majority of peaks lost and gained in IPF were intronic, with the second most commonly lost peak group residing in intergenic regions (Figure 2D), suggesting alterations in regulatory elements such as enhancers. Using the consensus peak sets, the genomic distribution of differentially accessible peaks was then compared (Figure 2E). Both lost and gained ATAC peaks in AT2-IPF samples were distributed throughout the genome, indicating profound epigenome-wide changes.

### Integrated transcriptomic and epigenomic analysis reveals *TRIP13* as a key target of epigenomic dysregulation in IPF

Significant differences in ATAC peak accessibility were calculated genome-wide using GenRich^(40)^. This revealed large shifts in the epigenome within AT2-IPF cells (Figure 3A). Two peaks stood out as the most significantly differentially accessible peaks between AT2-control and AT2-IPF samples: a lost peak located on chromosome 20 in an intergenic region between leucine-rich repeat neuronal (*LRRN4*) and FERM domain containing-kindlin 1 (*FERMT1*) genes (Chr20:6056852-6057730), and a gained peak within the distal tip of the p arm of chromosome 5 (Chr5:912228-913289), located in the 10^th^ intron of the thyroid receptor interactor 13 (*TRIP13)* gene.

**Figure 3:**
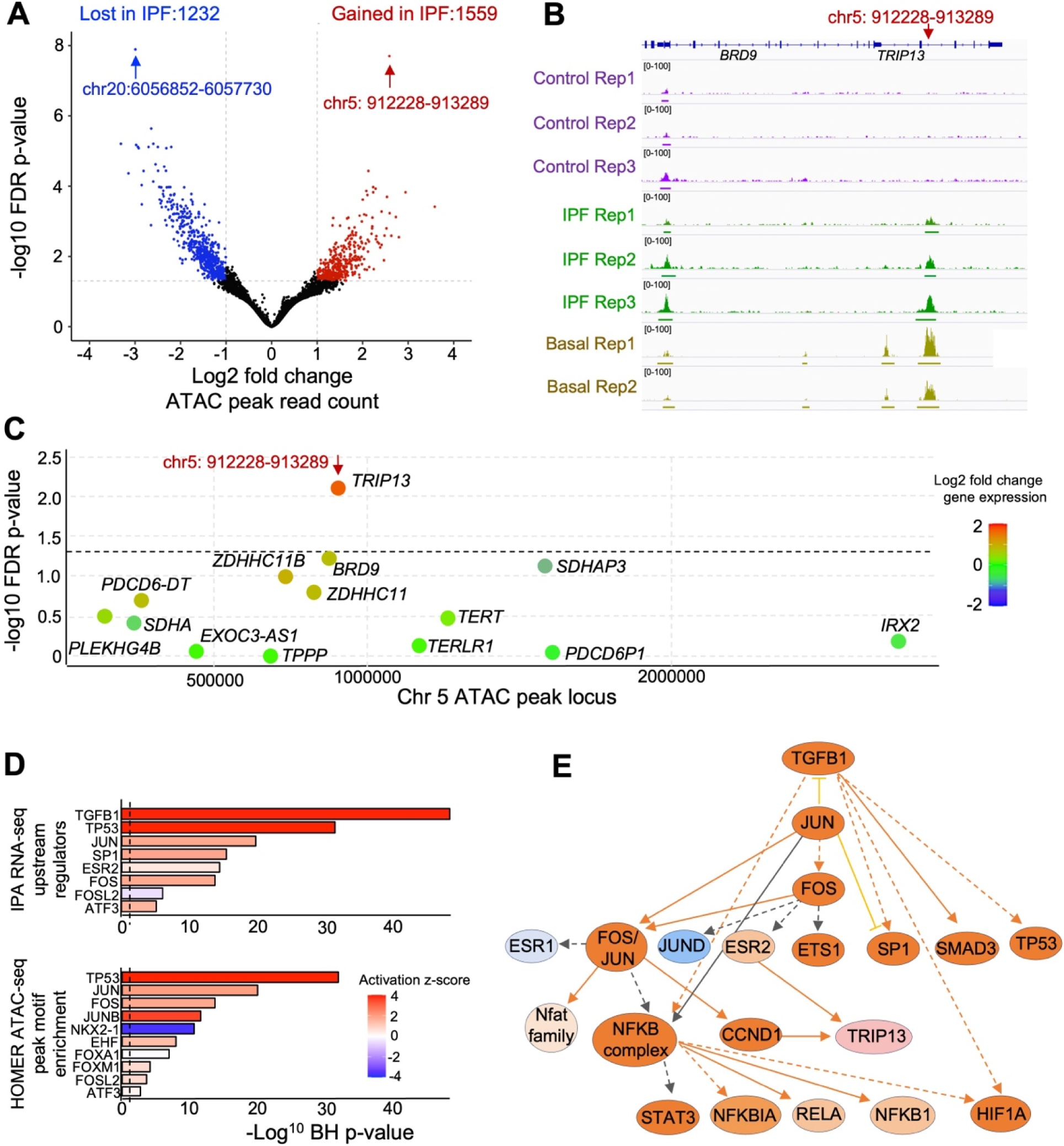
Integrated transcriptomic and epigenomic analysis reveals the *TRIP13* gene as a primary target of epigenomic dysregulation in IPF. **A.** Volcano plot of differential ATAC peak strength in AT2-control *vs.* AT2-IPF. X-axis = log2 fold change, y-axis = -log10 of the FDR-corrected p-value significance calculated in GenRich. Red = ATAC reads enriched in AT2-IPF consensus peak regions, blue = ATAC reads enriched in AT2-control consensus peak regions. Black = non-significant changes. Gray dotted lines = significance cutoffs. **B.** Read count enrichment and called peaks at the TRIP13 genomic locus displayed using Integrated Genomics Viewer (IGV). X-axis = Chromosome 5: 911711-913255 locus (hg38-based), Y-axis = peak height. AT2-control replicates = purple, AT2-IPF replicates = green, basal replicates = brown. **C.** Local Manhattan plot of the distal tip of the p-arm of chromosome 5. X-axis = genomic coordinates. Y-axis = -log of p-value for significance in gene expression changes. Dashed line indicates significance threshold. Each dot represents a known gene within the distal p arm of chromosome 5. Dot color reflects log2 fold change in gene expression between AT2-IPF and AT2-control. Red = elevated expression in AT2-IPF relative to AT2-control, blue = decreased expression in AT2-IPF relative to AT2-control. **D.** Bar plot of IPA of RNA gene expression differences in known upstream regulator transcription factors predicted to influence differential gene expression in AT2-IPF vs. AT2-control (top panel). Bar plot of HOMER-calculated transcription factor binding site (TFBS) motifs enriched in AT2-IPF vs. AT2-control consensus ATAC peaks r (bottom panel). Bars colors based on the IPA-calculated Z-sore of activation; red = predicted to be activated in AT2-IPF samples, blue = predicted to be inhibited in AT2-IPF samples. **E.** IPA network diagram of relationship between known upstream regulators predicted to be activated in AT2-IPF compared to AT2-control samples. Orange = predicted to be activated in AT2-IPF, blue = predicted to be inhibited in AT2-IPF.

We observed elevated levels of basal-associated genes in IPF AT2 cells (Figure 1). To determine if AT2-IPF cells were also acquiring ATAC changes associated with basal identity, we obtained ATAC-seq data from primary basal cells isolated from the tracheobronchial regions of the cartilaginous airways of non-IPF donors. We integrated these data (Figure S3) with AT2-control and AT2-IPF ATAC-seq data. GenRich^(40)^ was coupled with Diffbind^(41)^ to calculate read count and peak depth at each locus and visualize peak (dis)similarity as well as calculate significance of those differences (Table S3). We observed 88 sites of peak enrichment shared between AT2-IPF and basal identity (Figure S4, red points), suggesting limited adoption of basal-like peaks in AT2-IPF, rather than genome-wide acquisition of a basal identity. Interestingly, the AT2-IPF ATAC peak gained on chromosome 5 was the most highly significant peak both with respect to being gained in AT2-IPF and present in basal cells (Figure 3B).

The majority of enhancers are thought to target genes within ∼1 Mb^(42)^. We therefore integrated the transcriptomic data for the 2 Mb region on each side of the two top altered peaks from Figure 3A. *TRIP13* was the only significantly induced gene within the 2-Mb window of the Chr5:912228-913289 peak acquired in AT2-IPF cells (Figure 3C). Examining the 2-Mb region surrounding the top ATAC peak lost in AT2-IPF cells did not reveal a significantly downregulated gene (Figure S5). We therefore focused our molecular analyses on TRIP13 (see next section).

We used Ingenuity Pathways Analysis (IPA) to identify the top upstream regulators influencing the observed patterns of RNA disruption (Figure 3D, top) and Hypergeometric Optimization of Motif EnRichment (HOMER)^(39)^ to identify transcription factor binding sites (TFBS) enriched in AT2-IPF-specific ATAC peaks (Figure 3D, bottom). TGFB1, TP53 and members of the FOS/JUN families were identified as upstream regulators of RNA changes observed in AT2-IPF, while ATAC peaks in AT2-IPF were most enriched for TP53 and FOS/JUN family binding sites. Cross-referencing significantly enriched TFBS in HOMER with predicted upstream regulators of gene expression alterations of AT2-IPF in IPA revealed TGF-β as one of the top predicted significantly enriched upstream regulators in AT2-IPF-specific ATAC peaks, with known functional relationships to other activated upstream regulators such as FOS and JUN, and notably, TRIP13 (Figure 3E).

### TRIP13 is increased in IPF tissues and AT2 cells

Immunofluorescence (IF) staining showed increased TRIP13 expression in epithelial cells in IPF distal lung tissue compared to control (Figure 4A). In IPF, distinct subpopulations of AT2 cells, including cells expressing both alveolar and airway epithelial cells markers, have been identified in the alveolar compartment^(19,20,43,44)^. TRIP13 co-localized with alveolar type II (HTII-280) and basal (KRT5) epithelial cell markers (Figure 4A). Triple TRIP13^+^HTII-280^+^KRT5^+^ cells were also observed (Figure 4A, yellow arrows).

**Figure 4:**
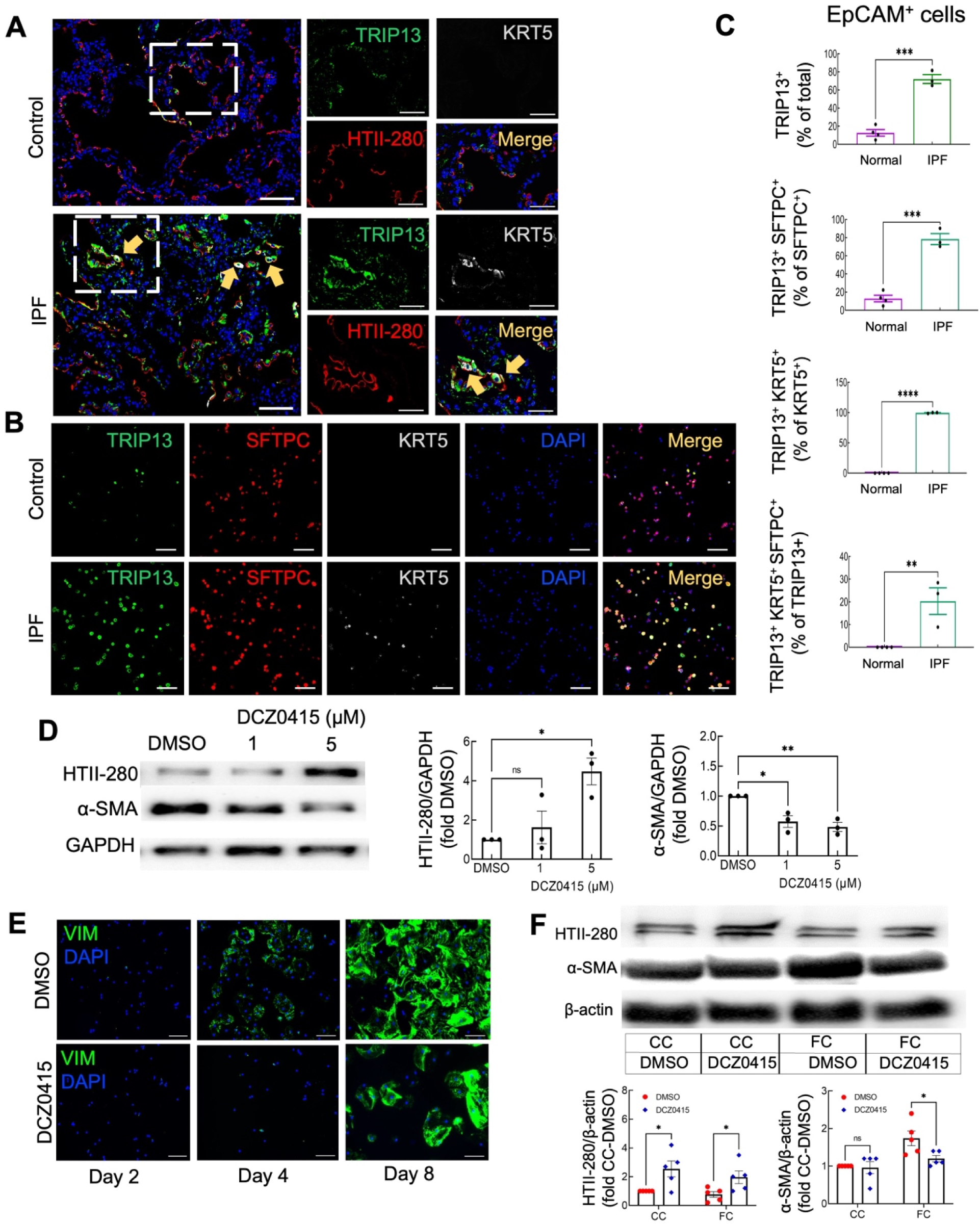
TRIP13 expression in control and IPF human lung and effect of TRIP13 inhibition on AT2 cells. **A.** Representative control (n = 4) and IPF (n = 4) lungs stained for TRIP13 (green), HTII-280 (red) and KRT5 (gray). Yellow arrows: TRIP13 co-expression with HTII-280 and KRT5. DAPI (blue): nuclear counterstain. Scale bars = 100 µm for lower magnification and 50 μm for higher magnification views (small squares). **B.** Representative cytospins of EpCAM^+^ cells isolated from control (n = 4) and IPF (n = 3) distal lung stained for TRIP13 (green), SFTPC (red) and KRT5 (gray). DAPI (blue): nuclear counterstain. Scale bars = 100 µm. **C.** Quantitative analysis of cytospins of EpCAM^+^ cells from B. Data are shown as mean ± SEM (* = P <0.05; ** = P < 0.01; *** = P < 0.001; **** = P < 0.0001). **D.** Representative western blots of AT2 cells cultured in 2D on plastic, treated with DMSO or DCZ0415 for 4 days (n=3) and probed for HTII-280 and α-SMA. Histograms at right show quantitation of western blot densitometric analyses. Data are shown as mean ± SEM (* = P < 0.05; ** = P ≤ 0.01). **E.** Representative images of AT2 cells cultured in 2D on chamber slides treated with DMSO or DCZ0415 (5 µM) for 2, 4, and 8 days, and stained for VIM (green) (n = 3). DAPI (blue): nuclear counterstain. Scale bars: 100 µm. **F.** Top panel: Representative western blots and quantitation of control lung tissue slice cultures treated with control cocktail (CC) or fibrotic cocktail (FC) along with DMSO or DCZ0415 (5 µM) for 2 days (n=5). Bottom panel: Quantitation of western blot densitometric analysis. Data are shown as mean ± SEM (* = P < 0.05).

Cytopsin preparations of EpCAM^+^ cells from AT2-IPF and AT2-control distal lung were co-stained for TRIP13, AT2 cell-specific surfactant protein C (SFTPC) and KRT5 (Figure 4B). TRIP13 was expressed in 72.1 ± 4.9% of IPF versus 12.8 ± 4.1% of control EpCAM^+^ cells (Figure 4C). Further classifying the EpCAM^+^ cell fraction as AT2 cells through dual EpCAM^+^SFTPC^+^ staining showed that 78.8 ± 6.1% of AT2-IPF cells were EpCAM^+^SFTPC^+^TRIP13^+^, whereas only 13.0 ± 3.8% of AT2-control cells were EpCAM^+^SFTPC^+^TRIP13^+^ (Figure 4C). Furthermore, 99.5 ± 0.5% of IPF EpCAM^+^KRT5^+^ cells expressed TRIP13. Of the IPF EpCAM^+^ cells that expressed TRIP13, 20.3 ± 5.9% co-expressed SFTPC and KRT5. None of the AT2-control cells expressed KRT5. Therefore, we observed a significant association between TRIP13 expression and the EpCAM^+^SFTPC^+^ cell population, while some of the latter had also acquired expression of basal-associated KRT5.

Upregulation of TRIP13 has been shown to promote EMT in cancer cells^(45–47)^ and IPF AT2 cells express higher levels of mesenchymal proteins^(48–50)^. We hypothesized that in IPF, TRIP13 promotes AT2 cell acquisition of mesenchymal features. To investigate this, we used a 2-dimensional (2D) cell culture model where control human HTII-280^+^ AT2 cells are grown on plastic, conditions which facilitate spontaneous EMT^(51)^. Cells were treated with the small molecule TRIP13 inhibitor, DCZ0415 (1 µM or 5 µM) or DMSO vehicle control. Under phase contrast light microscopy, DCZ0415-treated AT2 cell cultures exhibited mainly cuboidal (blue arrows) and fewer elongated (yellow arrows) cells (Figure S6), suggesting maintenance of an epithelial phenotype. DCZ0415 treatment for 4 days partially prevented EMT and promoted AT2 cell maintenance as shown by significantly reduced fibrotic marker α-smooth muscle actin (α-SMA) and increased HTII-280 expression, respectively (Figure 4D). However, higher concentrations (10 µM) of DCZ0415 caused cell death. To serially examine TRIP13 inhibitory effects on acquisition of mesenchymal features in HTII-280^+^ AT2 cells in 2D culture, we evaluated expression of vimentin (VIM) on days 2, 4, and 8 by IF staining in the presence or absence of DCZ0415 (5 µM). DCZ0415 significantly reduced VIM protein expression and its onset was delayed compared to untreated cells (Figure 4E, Figure S6).

Treatment of short-term human lung tissue slice cultures with profibrotic and proinflammatory agents mimics some features of IPF^(52–54)^. To further explore the potential antifibrogenic effect of DCZ0415, we treated control human lung tissue slices with a fibrotic cocktail (FC) consisting of 5 ng/ml TGF-β, 10 ng/ml platelet-derived growth factor-AB (PDGF-AB), 10 ng/ml tumor necrosis factor-alpha (TNF-α) and 5 μM lysophosphatidic acid (LPA) in the presence or absence 5 µM DCZ0415. FC promotes EMT and has also been suggested to promote the transdifferentiation of AT2 cells to airway-like epithelial cells^(55)^. DCZ0415 maintained the AT2 cell phenotype as shown by increased HTII-280 expression in the presence of FC (Figure 4F). DCZ0415 also significantly blocked FC-stimulation of α-SMA, indicating the ability to inhibit EMT (Figure 4F) and suggesting that epigenomic activation of *TRIP13* in AT2-IPF may contribute to an EMT phenotype in IPF.

## DISCUSSION

This study demonstrates profound genome-wide transcriptomic and epigenomic alterations in AT2-IPF cells, enabling them to adopt hallmarks of EMT. Among others, AT2-IPF cells had key ATAC peaks in common with basal cells. The top ATAC peak gained in AT2-IPF occurred within intron 10 of *TRIP13*. *TRIP13* expression is significantly elevated at the mRNA and protein level in AT2-IPF compared to AT2-control cells. Using the TRIP13-selective small molecule inhibitor, DCZ0415, we show that blocking TRIP13 mitigates AT2 cell EMT. Together these observations suggest that TRIP13 contributes functionally to hallmarks of IPF.

TRIP13 is a nuclear protein that interacts with the hormone-dependent transcription factor family and plays a role in DNA repair^(56)^. TRIP13 is implicated in EMT in lung cancer through modulation of TGF-β signaling^(47, 57–59)^. TRIP13 inhibition blocks EMT in colorectal cancer^(46)^ and multiple myeloma^(60)^. TRIP13 modulates TGF-β-SMAD signaling in multiple disease states^(57, 58)^, and is itself a target of TGF-β signaling in mice^(63)^. This study and others have found elevated TGF-β signaling and activation of downstream targets in IPF^(64–67)^. TGF-β was the top upregulated upstream regulator in IPF-AT2 cells. Indeed, TGF-β signaling has been linked to the induction of transitional cell states^(21, 68)^ implicated in IPF pathogenesis. DCZ0415 is primarily being pursued as an anti-cancer agent; however, our study suggests that DCZ0415 may also be effective against fibrotic diseases.

SMAD signaling is the major canonical nuclear target of TGF-β signaling, and dysregulated SMAD signaling is implicated as a critical driver of IPF^(16, 69–71)^. A major component of SMAD function is recruitment of epigenomic modulators to influence chromatin structure^(72,73)^. Altered SMAD signaling influences recruitment and distribution of histone acetylase complexes genome-wide in multiple cellular models^(73,74)^, and may partially explain the epigenomic changes observed in this study.

Several additional transcription factors were implicated in alterations to the epigenomic landscape, including prominently, members of the AP-1 (FOS/JUN) family. AT2 cells can undergo pro-fibrotic cell fate transitions^(75)^, which may be enhanced by activated AP-1 family member signaling^(76)^. TP53 was also identified as a major upstream regulator predicted to be heavily enriched within open chromatin regions in IPF. Ample evidence supports a central role for TP53 signaling in IPF through dysregulated injury resolution^(19, 77–79)^. Additional studies on these transcription factors and regulatory networks promise to yield further critical insights into disease pathogenesis.

In summary, we demonstrate that IPF AT2 cells undergo widespread epigenomic deregulation and activation of mesenchymal and airway genes. The most prominent ATAC peak gained was in *TRIP13*. Notably, at the transcriptional level, *TRIP13* is not the most significantly induced gene in AT2-IPF, and in the absence of epigenomic profiling its significance may have been overlooked, underscoring the value of integrated multiomic analysis. Suppression of fibrosis in 2D and lung slice cultures by TRIP13 inhibition points to the key role this gene may play in epithelial plasticity in IPF, supporting further investigation of DCZ0415 as a potential IPF treatment.

### STAR METHODS

#### KEY RESOURCES TABLE

| REAGENT OR RESOURCE | SOURCE | IDENTIFIER |
| --- | --- | --- |
| **Antibodies** |  |  |
| Chicken IgY anti-human KRT5 polyclonal, unconjugated; 1:500 IF | Biolegend | Cat# 905904<br>RRID: AB_2721742 |
| Mouse IgM anti-human HTII-280 monoclonal, unconjugated; 1:200 IF, 1:500 WB, 1:50 cytospin | Terrace Biotech | Cat# TB-27AHT2-280<br>RRID: AB_2832931 |
| Rabbit IgG anti-human TRIP13 polyclonal, unconjugated; 1:100 | Proteintech | Cat# 19602-1-AP<br>RRID: AB_10642702 |
| Mouse IgG anti-human SFTPC monoclonal, unconjugated; 1:50 IF, 1:100 cytospin | Santa Cruz Biotech | Cat# sc-518029<br>RRID: AB_2937075 |
| Rabbit IgG anti-VIM monoclonal, unconjugated; 1:100 IF | Cell Signaling | Cat# 5741S<br>RRID: AB_10695459 |
| Mouse IgG anti-human CDH1 monoclonal, unconjugated; 1:100 | Cell Signaling | Cat# 14472S<br>RRID: AB_2728770 |
| Goat anti-chicken polyclonal, Alexa flour 488- conjugated; 1:200 | Invitrogen | Cat# A-11039<br>RRID: AB_2534096 |
| Goat anti-mouse polyclonal, Alexa flour 568- conjugated; 1:200 | Invitrogen | Cat# A-21043<br>RRID: AB_2535712 |
| Goat anti-mouse polyclonal, biotin-conjugated; 1:200 | Vector Laboratories | Cat# BA-9200-1.5<br>RRID: AB_2336171 |
| Goat anti-rabbit polyclonal, biotin-conjugated; 1:200 | Vector Laboratories | Cat# BA-1000-1.5<br>RRID: AB_2313606 |
| Mouse IgG anti-human ACTA2 monoclonal; 1:1000 | Sigma Aldrich | Cat# A5228<br>RRID: AB_262054 |
| Mouse IgG anti-human ACTB monoclonal, unconjugated; 1:500 | Santa Cruz | Cat# sc-47778<br>RRID: AB_626632 |
| Rabbit IgG anti-human GAPDH monoclonal, unconjugated; 1:1000 | Cell Signaling | Cat#2118S<br>RRID: AB_561053 |
| Horse anti-mouse polyclonal, HRP-conjugated; 1:2000 | Cell Signaling | Cat# 7076S<br>RRID: AB_330924 |
| Goat anti-rabbit polyclonal, HRP-conjugated; 1:2000 | Cell Signaling | Cat# 7074S<br>RRID: AB_2099233 |
| Mouse IgG<sub>1</sub> anti-human SCGB1A1 monoclonal, unconjugated; 1:100 | Santa Cruz Biotech | Cat# sc-365992<br>RRID: AB_10915481 |
| Rabbit IgG anti-human CD45 polyclonal, unconjugated; 1:200 | Santa Cruz Biotech | Cat# sc-25590<br>RRID: AB_2174143 |
| Mouse IgG2b anti-human acetylated tubulin monoclonal, unconjugated; 1:200 | Millipore Sigma | Cat# T6793<br>RRID: AB_477585 |
| Rabbit IgG anti-human KRT5 monoclonal, unconjugated; 1:100 cytospin | Abcam | Cat# Ab52635<br>RRID: AB_869890 |
| Mouse IgM anti-human VIM monoclonal, unconjugated; 1:200 | Sigma-Aldrich | Cat# V2258<br>RRID: AB_2315587 |
| Rabbit IgG anti-human MUC5B polyclonal, unconjugated; 1:100 | Santa Cruz Biotech | Cat# sc-20119<br>RRID: AB_2282256 |
| Mouse IgG1 anti-human TTF1 (NKX2-1) monoclonal; unconjugated 1:100 | Novocastra | Cat# TTF1-1-LCE |
| Rabbit anti-human SFTPB polyclonal, unconjugated; 1:100 | Millipore Sigma | Cat# AB3430<br>RRID: AB_11210466 |
| IgG from human serum | Sigma Aldrich | Cat# I4506<br>RRID: AB_1163606 |
| Mouse IgG1 anti-human CD326 (EpCAM)-conjugated microbeads | Miltenyi Biotec | Cat# 130-061-101<br>RRID: AB_2832928 |
| Anti-mouse IgM -conjugated magnetic beads | Miltenyi Biotec | Cat# 130-047-302<br>RRID: AB_244359 |
| **Chemicals, Peptides, and recombinant proteins** |  |  |
| Streptavidin, Alexafluor 647-conjugate; 1:200 | Invitrogen | Cat# S32357 |
| Vectashield Antifade Mounting Medium with Propidium Iodide (PI) | Vector Laboratories | Cat# H-1300-10 |
| Transforming growth factor-β | PeproTech | Cat# 100-21 |
| Platelet-derived growth factor-AB | Thermo Fisher | Cat# PHG0134 |
| Tumor necrosis factor-α | Bio-Techne | Cat# #210-TA-020 |
| Penicillin-streptomycin | Sigma-Aldrich | Cat# P0781 |
| DME-F12 | Sigma | Cat# D64421 |
| DMEM High Glucose | GIBCO | Cat# 21063 |
| Gentamycin | Sigma | Cat# G1272 |
| Fungizone | Sigma | Cat# A2411 |
| Hank's Balanced Salt Solution 1x media without $\text{Ca}^{2+}$ or $\text{Mg}^{2+}$ | VWR | Cat #21-021-CV |
| Elastase | Worthington | Cat# LS002280 |
| Percoll | Sigma | Cat# P1644 |
| FcR Blocking Reagent | Miltenyi Biotec | Cat# 130-059-901 |
| Trypan blue | Sigma | Cat# T8154 |
| Paraformaldehyde |  |  |
| Primocin | InvivoGen | Cat# NC9141851 |
| Protease XIV | Sigma | Cat# P5147 |
| DNase | Sigma | Cat# DN25 |
| DMEM/F12 | Thermo Fisher | Cat# 11330032 |
| ACK lysis buffer | Thermo Fisher | Cat# #A1049201 |
| Accutase | Innovative Cell Technologies | Cat# AT104 |
| Purecol® | Advanced BioMatrix | Cat# 5005 |
| Airway epithelial growth media | Promocell | Cat# C-21160 |
| CAS block | Thermo Fisher | Cat# 008120 |
| Vectashield Antifade Mounting Medium with 4',6-diamidino-2-phenylindole (DAPI) | Vector Laboratories | Cat# H-1200-10 |
| Amphotericin B | Sigma-Aldrich | Cat# A2411 |
| Dispase | Corning | Cat# 354235 |
| DNase I | Roche | Cat# 10104159001 |
| Fetal Bovine Serum | GenClon | Cat# 25-514 |
| Bovine type I collagen | Advanced BioMatrix | Cat# 5005 |
| lysophosphatidic acid (LPA) (#62215, Cayman Chemical,) | Cayman Chemical | Cat#62215 |
| DCZ0415, TRIP13 inhibitor | MedChemExpress | Cat# HY-130603 |
| Protease inhibitor cocktail | Sigma-Aldrich | Cat# P1860 |
| Red blood cell lysis buffer | Roche | Cat# 11814389001 |
| Citrate-based antigen retrieval | Vector Laboratories | Cat# H-3300-250 |
| TrueBlack lipofuscin autofluorescence quencher | Biotium | Cat# 23007 |
| Phosphatase inhibitor cocktail 2 | Sigma-Aldrich | Cat# P5726 |
| Polyvinylidene difluoride (PVDF) membranes | Bio-Rad | Cat# #1620177 |
| ECL Substrate | Thermo Fisher | Cat# 34580 |
| Critical commercial assays |  |  |
| LS columns | Miltenyi Biotec<br>Cat#130-042-401 |  |
| EZ single cytofunnels | Thermo Fisher | Cat# 1439045 |
| Aurum Total RNA Mini kit | BioRad | Cat# 7326820 |
| ATAC-seq kit | Active Motif | Cat# 53150 |
| Deposited data |  |  |
| Data files for bulk RNA-seq and ATAC-seq in AT2 cells | This paper | GSE245965 |
| Experimental models: plasmids |  |  |
| pTXB1-Tn5ES |  | Reference: (80) |
| Oligonucleotides |  |  |
| Nextera i7 dual indexing primers |  |  |
| Software and Algorithms |  |  |
| BioVenn generator |  | Reference: (81) |
| DESeq2 (v1.42.0) | Love, Michael I et al. | Reference: (32) |
| FASTQC |  |  |
| Ggplots2 |  |  |
| HOMER (v4.11, 10-24-2019) | Heinz, Sven et al. | http://homer.ucsd.edu/homer/index.html |
| IGV (v2.16.2) | Robinson, James T et al. | https://igv.org/ |
| IPA (v101138820) | QIAGEN | https://digitalinsights.qiagen.com/products-overview/discovery-insights-portfolio/analysis-and-visualization/qiagen-ipa/ |
| PANTHER |  | Reference: (82) |
| R 4.2.3 | R | https://www.r-project.org/ |
| R Studio (v2023.06.0+421) | Rstudio | https://www.rstudio.com |
| STAR aligner |  | Reference: (83) |
| ImageJ software |  | Reference: (84) |
| Trimmomatic |  | Reference: (85) |
| Other |  |  |
| Falcon cells | VWR | Cat# 734-0004 |
| Shandon CytoSpin III cytocentrifuge | GMI | SKU: 8358-30-0001 |
| #10 scalpel | Exel International | Cat# 29550 |
| Leica DMI8 Fluorescent microscope | Leica Microsystems |  |
| Leica TCS SP8 confocal microscope | Leica Microsystems |  |
| Slide-a-Lyzer dialysis cassette | Thermo Fisher | Cat# 6630 |
| Amicon Ultracel 50K centrifugal Filter | Millipore Sigma | #UFC505008 |
| MultiTherm shaker | Benchmark Scientific | Cat# H5000-H |
| Agilent 4200 Tape Station |  |  |
| Gauze mesh strainer | Corning | Cat# 352360 |
| 40 µM strainer | Corning | Cat# 352340 |
| HistoCore Pegasus tissue processor | Leica Biosystems |  |
| HistoStar Embedding Center | Epredia |  |
| Keyence fluorescence microscope BZ-X800 | Keyence Corporation |  |
| ChemiDoc™ MP imaging system | Bio-Rad |  |

#### Resource availability

##### Lead contact

Further information and requests for resources and reagents should be directed to and will be fulfilled by the lead contact, Zea Borok.

##### Materials availability

No newly generated materials are associated with the paper.

##### Data and code

*All data reported in this paper will be shared by the lead contact upon request*.

- This paper does not report original code.
- Any additional information required to reanalyze the data reported in this paper is available from the lead contact upon request.

## Method Details

### Human lung tissue acquisition

IPF tissues were obtained from explanted lungs from consented patients undergoing lung transplant with the approval of Institutional Review Boards (IRB) at USC (IRB# HS-14-00829 and IRB# HS-18-00273) and UCSD (IRB# 210575 and IRB# 201252). Deidentified non-IPF (control) lungs from deceased donors were obtained under material transfer agreements with the International Institute for the Advancement of Medicine (IIAM) and the University of North Carolina Marsico Lung Institute. IPF and control human lung tissues were either freshly processed for isolation of epithelial cells (for RNA-seq and ATAC-seq) or histologic evaluation, or cryopreserved in freezing media for later use (for cell and lung slice cultures).

### Lung tissue preparation for histology

For all lungs, small pieces of peripheral tissue were fixed in 4% paraformaldehyde (PFA) overnight at 4°C. After washing with 1X phosphate buffered saline (PBS), fixed tissues were stored in 70% ethanol at 4°C until paraffin embedding. Fixed tissues were processed through a HistoCore Pegasus tissue processor (Leica Biosystems) and then paraffin embedded using the HistoStar Embedding Center (Epredia).

### Human lung tissue cryopreservation

Lung tissues were cut into small pieces (∼4 mm^3^) and washed with Dulbecco’s Modified Eagle Medium/Nutrient Mixture F-12 (DMEM/F-12) containing 1% penicillin-streptomycin (#P0781, Sigma-Aldrich) and 1% amphotericin B (#A2411, Sigma-Aldrich). Tissues were transferred into 2 mL or 5 mL cryovials containing 1 mL or 2.5 mL freezing media comprised of 90% fetal bovine serum (FBS) and 10% dimethyl sulfoxide (DMSO) or FrostaLife Cryopreservation Solution (#LM-0015, LifeLine Cell Technology). Cryovials were transferred into freezing containers and kept at -80°C overnight and subsequently transferred into liquid nitrogen tanks for long-term storage.

### AT2 cell isolation from fresh human lungs for RNA-seq and ATAC-seq

Lung parenchyma was macro-dissected to a depth of ∼2 mm^3^ to isolate alveolar tissue. The visceral pleura was removed using forceps and alveolar tissue was then minced into fine pieces with scissors. Tissue was then washed in 50:50 media, which is a 50:50 mixture of DMEM/F-12 (#D64421, Sigma) and DMEM High Glucose (#21063, GIBCO) supplemented with 1% penicillin/streptomycin (#P-0781, Sigma-Aldrich) and 1% Fungizone (A2411, Sigma-Aldrich). EpCAM^+^ cells were isolated as previously reported for samples used for RNA-seq and ATAC-seq^(29)^. Following overnight incubation in DMEM/F-12 containing 1X dispase (#354235, Corning, NY) at 4°C, the tissue was again minced briefly using scissors and resuspended in Hank’s Balanced Salt Solution (HBSS) without calcium or magnesium (#21-021-CV, VWR) containing 13 U/mL of elastase (#LS002280, Worthington Biochemical Corp) and incubated for 30 minutes at 37°C. Tissue homogenate was serially filtered through gauze mesh and 100 µm (#352360, Corning) and 40 µm (#352340, Corning) cell strainers, then pelleted and resuspended in 50:50 media. After purification of epithelial cells using Percoll (#P1644, Sigma) density gradient centrifugation, macrophages and leukocytes were further depleted by IgG panning by adherence to human IgG-coated (#I4506, Sigma-Aldrich) Petri dishes. After FcR blocking (#130-059-901, Miltenyi Biotec), magnetic-activated cell sorting (MACS) was performed using human CD326 (EpCAM) microbeads (#130-061-101, Miltenyi Biotec) with LS columns (#130-042-401, Miltenyi Biotec) according to the manufacturer’s instructions. Trypan blue (#T8154, Sigma) was used to assess cell viability. For preparation of cytospin slides for evaluation of AT2 cell purity and immunostaining, cells were fixed in 4% PFA for 10 minutes, washed with PBS and resuspended in PBS. Cytospins were prepared using EZ single cytofunnels (#1439045, Thermo Fisher Scientific) and a Shandon CytoSpin III cytocentrifuge (GMI). Cytospin slides were stored at -20°C until use.

### Alveolar epithelial cell isolation from cryopreserved lung tissues

Cryopreserved IPF and control human lung tissues were thawed rapidly in a 37°C water bath. Under sterile conditions, tissues were washed with DMEM/F-12 containing 1% penicillin-streptomycin (#P0781, Sigma-Aldrich) and 1% amphotericin B (#A2411, Sigma-Aldrich) and minced with razor blades in Petri dishes. Minced tissues were digested with an enzyme cocktail containing 1X dispase (#354235, Corning), 13 U/mL elastase (#LS002280, Worthington Biochemical Corp) and 0.33 U/mL DNase I (#10104159001, Roche) in DMEM/F-12 for 1h at 37°C in a shaking water bath. Enzymatic activity was stopped with 10% FBS and then serial filtration of the tissue homogenate was performed as described above for cell isolation from fresh lung tissues and previously reported (Marconett 2013). Red blood cells were lysed using red blood cell lysis buffer (# 11814389001, Roche). The remaining steps, including macrophage and leukocyte depletion using IgG panning and MACS for selection of EpCAM^+^ cells, were performed as described above for cell isolation from fresh lung tissues. Further enrichment of AT2 cells from control human lung used for cell culture was performed with MACS using HTII-280 monoclonal mouse IgM antibody (1:50) (#TB-27AHT2-280, Terrace Biotech) along with magnetic bead-labeled anti-mouse IgM antibody (#130-047-302, Miltenyi Biotec) with LS columns. Cytospins were processed as described above.

### Isolation of basal cells from non-IPF donors

Isolation of primary human airway epithelial cells (HAECs) was performed as previously described^(86)^. Briefly, proximal airways from the trachea, main stem bronchi and 2-3 further branching generations of the cartilaginous airways were dissected into sections between 1-4 cm^2^ in size and placed in a solution (%w/v) of 0.1% Protease XIV (#P5147, Sigma) and 0.001% DNase (#DN25, Sigma) in DMEM/F12 (#11330032, ThermoFisher) overnight at 4°C. Using a #10 scalpel (#29550, Exel International), epithelial tissue was gently scraped off, collected in DMEM/F12, and centrifuged at 400 x g for 5 minutes. After red blood cell lysis in ACK lysis buffer (#A1049201, ThermoFisher), epithelial cells were dissociated to single cells by incubation with Accutase (#AT104, Innovative Cell Technologies) at 37°C for 30 minutes.

### RNA and ATAC material extraction and library preparation

For RNA preparation, 6 x 10^6^ AT2 cells isolated from fresh lung tissues were pelleted and stored at -80°C. RNA was extracted using the Aurum Total RNA Mini kit (# 7326820, BioRad) according to manufacturer guidelines. 2 µg of total RNA then underwent bulk RNA library preparation with PolyA selection performed by Genewiz (now Azenta). Samples were barcoded using Illumina adapters to allow for multiplexed sequencing. For ATAC of AT2 cells, the ATAC-seq assay was performed as described^(87)^ with two modifications: nuclei were transposed using a purified Tn5 isolated from bacterial isolations of pTXB1-Tn5ES clones as previously described^(80)^, with substitution of tubing for a Slide-a-Lyzer dialysis cassette (#6630, ThermoFisher) and an Amicon Ultracel 50K centrifugal Filter (#UFC505008, Millipore Sigma) to increase protein yield. Library fragments were amplified using NEB Next Ultra II Q5 Master Mix. For ATAC of basal cells, 5-50 X 10^3^ cells isolated from fresh lung tissues were placed in 0.1 to 0.25 mL of ATAC lysis buffer (10 mM Tris-HCl, pH 7.5, 10 mM NaCl, 3 mM MgCl_2_), dependent on cell number, and immediately processed for ATAC library generation using the Active Motif ATAC-seq kit (#53150, Carlsbad, CA) according to manufacturer guidelines, including Nextera i7 dual indexing primers for library construction. Integration of indexing primers was performed using either the standard Illumina Tn5, per manufacturer’s protocol, or with Tn5 isolated from bacterial isolations of pTXB1-Tn5ES clones as previously described^(80)^ with substitution of tubing for a Slide-a-Lyzer dialysis cassette (#6630, ThermoFisher) and an Amicon Ultracel 50K centrifugal Filter (#UFC505008, Millipore Sigma) to increase protein yield. A H5000-H MultiTherm shaker (Benchmark Scientific) was used to ensure uniformity in library construction. The Agilent 4200 Tape Station system was used for assessment of ATAC fragment quality throughout library construction.

### Next generation sequencing

Bulk RNA paired end sequencing and bulk ATAC paired end sequencing was performed at Genewiz, using an Illumina HiSeq at 2x150 bp. RNA-seq was performed using single index chemistry, whereas ATAC-seq was performed using dual-index chemistry.

### Bulk AT2 RNA-seq analysis

Raw FASTQ files underwent quality control analysis using FASTQC. Adapter sequences were then trimmed from raw FASTQ reads using Trimmomatic^(85)^. Reads were then filtered to remove those of low quality. Retained reads had a quality score of >30 per base for at least 90% of the read. Read metrics are contained in Figure S1. Reads were then aligned to the Hg38 genome with STAR aligner^(83)^ using the Ensembl v110 (July 2023 release) transcriptome build. Counts per gene per sample were then used for calculating differential expression analysis in DeSeq2^(32)^ in R (version 1.42.0). Sex distribution in samples was corrected for by removing genes on the X and Y chromosomes. Heatmaps displaying differentially expressed genes were visualized using the pheatmap functionality in R. FDR-corrected p-values were imported into Ingenuity Pathway Analysis^(88)^ from Qiagen. PANTHER^(82)^ was used as a secondary means of biological processes enrichment, performed using the statistical overrepresentation test^(89)^. Volcano plots of gene expression levels were generated using ggplots2 in R.Bulk.

### Bulk AT2 and basal ATAC-seq analysis

Preprocessing of raw ATAC-seq data to generate clean read files was performed as for the RNA samples. Cleaned ATAC-seq reads were aligned to the hg38 genome using bwa aligner^(90)^. Peak enrichment was subsequently calculated using the GenRich peak caller^(40)^ using the intrinsic background of the sample. Peaks were then annotated to nearest-neighbor genes using the annotatepeaks.pl function in HOMER^(39)^, which also binned peaks into categories relative to the genomic structures in which they occurred in (e.g., promoter, enhancer, etc). Heatmaps were generated by utilizing the annotatepeaks.pl output from HOMER and sorting based on peak intensity. FindMotifsGenome.pl in HOMER was used to calculate the significance of enrichment of transcription factor binding sites (TFBS) within peak regions, using the opposite sample type as the background file to account for open region composition. Venn diagrams of peak overlaps were generated using the BioVenn generator^(81)^. Peak and coverage tracks were visualized using BAM alignment files loaded into the Integrated Genomics Viewer (IGV)^(91)^.

### Data Accessibility

Bulk RNA-seq and ATAC-seq data from AT2-control, AT2-IPF, and basal cells presented in this study are deposited in the Gene Expression Omnibus (GEO) under record GSE245965.

### 2D culture of AT2 cells for inhibitor studies

Control human HTII-280^+^ AT2 cells isolated from cryopreserved lung tissue were cultured at 37°C under humidified conditions in 5% CO_2_ in DMEM/F-12 supplemented with 5% heat inactivated FBS (#25-514, GenClon) and 100 μg/ml primocin (#NC9141851, InvivoGen) in 8-well chamber slides (for immunofluorescence (IF)) and 24-well cell culture plates (for western blot analysis) precoated with bovine type I collagen (# 5005, Advanced BioMatrix). Where indicated, starting from day zero (D0), cells were treated with either 1 µM or 5 µM of TRIP13 inhibitor, DCZ0415 (#HY-130603, MedChemExpress). Media with DCZ0415 or vehicle (DMSO) were changed every other day. On days 2, 4 and 8 in culture, cells were fixed in 4% PFA for 10 minutes, washed with PBS and stored at -80°C in PBS containing 1% BSA for IF staining. For western blot analysis cells were lysed on day 4 following treatment.

### Lung tissue culture

Lung tissues were cultured as previously described with minor modifications^(92)^. After thawing rapidly in a 37°C water bath, lung tissues were filled with 2% low melt agarose kept at 37°C using a syringe pump. Agarose-tissue blocks were sliced into smaller pieces (∼1 mm^3^) using a razor blade. Tissue slices were cultured for two days in DMEM/F-12 containing 100 μg/ml primocin at 37°C under humidified conditions in 5% CO_2_ in 24-well plates (one slice per well) under submerged conditions with daily medium changes. Starting from Day 0, tissue slices were treated with FC^(54)^ consisting of 5 ng/ml TGF-β (#100-21, PeproTech), 10 ng/ml PDGF-AB (#PHG0134, Thermo Fisher), 10 ng/ml TNF-α (#210-TA-020, Bio-Techne) and 5 μM LPA (#62215, Cayman Chemical), or a control cocktail (CC) composed of water, PBS, and 1 uM acetic acid. FC- or CC-treated cells were then treated with either 5 μM DCZ0415 or DMSO vehicle as indicated. Lung slices were lysed on day 2 and stored at -20°C for western analysis.

### IF staining of cytospins, lung tissues and cultured AT2 cells

IF staining was performed as previously described with minor modifications^(93)^. For IF staining of cytospin preparations, slides were first allowed to reach room temperature and were then blocked with CAS block (#008120, Invitrogen, Waltham, MA) at room temperature for 30 minutes to 1 hour. For lung tissue staining, tissue sections (5 μm) after being baked at 56 °C for 1 hour, were dewaxed in xylene and rehydrated through graded ethanol. Following microwave citrate-based antigen retrieval (#H-3300-250, Vector Laboratories, Newark, CA), tissue sections were treated with TrueBlack lipofuscin autofluorescence quencher (#23007, Biotium, Fremont, CA) for 2 minutes. Sections were blocked in CAS block solution (#008120, Invitrogen) containing 5% normal goat serum for 1 hour at room temperature. For staining of AT2 cells cultured in chamber slides, samples were thawed at room temperature and then permeabilized for 10 minutes at room temperature with 0.2% Triton X-100 in PBS. Cytospins, lung sections and cultured cells were then incubated with primary antibodies in blocking buffer (listed in the Key resource table) at 4°C overnight. Slides were washed with PBS three times and then incubated with respective Alexa Fluor conjugated or biotinylated secondary antibodies in blocking buffer for 1 hour at room temperature. After washing with PBS three times, slides incubated with biotinylated secondaries were incubated with Alexa Fluor conjugated streptavidin in blocking buffer for 30 minutes at room temperature. After washing with PBS three times, slides were mounted with media containing 4′,6-diamidino-2-phenylindole (DAPI). Fluorescent images were captured at the same settings for all slides within an experimental group (Leica DMI8, Leica Microsystems, Buffalo Grove, IL and Keyence BZ-X800, Keyence Corporation of America, Itasca, IL). For cell counting, five random images were taken using the 20X lens of a Leica DMI8 fluorescence microscope and cell counting was performed using ImageJ software^(84)^. A minimum of 600 cells (stained for DAPI) was counted per cytospin slide. Non-overlapping visual fields of IF stained cultured cells were taken (minimum six images per well) using the 20X lens of a BZ-X800 Keyence fluorescence microscope and quantification of the areas of cytoplasmic and nuclear staining was automatically performed in each captured image using the hybrid cell count software, BZ-X analyzer (Keyence Corporation).

### Western blotting

To prepare lysates from cells and lung slices, samples were washed twice in ice-cold PBS and incubated on ice in lysis buffer containing 100 mM sodium chloride, 10 mM Tris-HCl (pH 7.5), 2 mM ethylene-diamine-tetra-acetic acid (EDTA), 0.5% (w/v) deoxycholate, 1% (v/v) Triton-X, 1% (v/v) protease inhibitor cocktail (#P1860, Sigma-Aldrich), 1% (v/v) phosphatase inhibitor cocktail 2 (#P5726, Sigma-Aldrich) and 1 mM magnesium chloride as previously described^(94)^. Adherent cells were detached using a cell scraper and lung slices were minced with scissors. Lysates were transferred to 1.7 mL Eppendorf tubes and centrifuged at 11,000 g for 10 min at 4°C after which supernatants were aspirated and stored at -20°C until required. Protein samples (usually 30 μg) were reduced by adding sample buffer (10 μL) containing 0.125 M Tris-HCL (pH 6.8), 20% (v/v) glycerol, 4% (w/v) SDS, 0.1% (w/v) bromophenol blue, 1% (v/v), 10% (v/v) β-mercaptoethanol and heating to 95°C for 5 minutes. Equal amounts of protein were resolved using 10% sodium dodecyl sulfate polyacrylamide gel electrophoresis (SDS-PAGE) at 200 V for 45 minutes. Protein was electrophoretically transferred to Immuno-blot polyvinylidene difluoride (PVDF) membranes (#1620177, Bio-Rad, Hercules, CA) in transfer buffer at 100 V for 1 hour on ice. Membranes were blocked with 5% (w/v) skim milk in Tris-buffered saline with 0.1% Tween 20 (TBS-T) for 1 hour. After a 5-minute wash with TBS-T, membranes were incubated with primary antibodies (Key Resources Table) diluted with 3% (w/v) bovine serum albumin (BSA) in TBS-T at 4°C overnight. The following day, membranes were washed three times (each for 5 minutes) with TBS-T and then incubated with horseradish peroxidase (HRP)-conjugated appropriate secondary antibodies diluted in 5% milk in TBS-T at room temperature for two hours. Finally, after three additional washes, the protein of interest was detected using ECL Substrate (cat. no. 34580, ThermoFisher) and imaged using a ChemiDoc™ MP imaging system (Bio-Rad, Hercules, CA). The levels of glyceraldehyde 3-phosphate dehydrogenase (GAPDH) protein or beta actin were used as controls for loading. Densitometric analysis of the bands was carried out using ImageJ^(84)^.

### Statistical Analysis

RNA-seq expression and ATAC-seq peak occupancy comparisons were performed using DEseq2 or GenRich, respectively, with genome-wide false discovery rate corrections applied. Likewise, Ingenuity Pathways Analysis was corrected for overrepresentation using the Benjamini-Hochberg method. Statistical analysis for cytospins, lung slices, and cultured cells were conducted using GraphPad Prism 9.0 software. Data were analyzed as grouped data and presented in figures as mean ± SEM from “n” donors. For one-factor analysis one-way ANOVA or unpaired t-test and for two-factor analysis two-way ANOVA were used with Bonferroni’s post hoc test correction for multiple comparisons. P ≤ 0.05 were considered to be statistically significant (* = P ≤ 0.05; ** = P ≤ 0.01; *** = P ≤ 0.001; **** = P ≤ 0.0001).

**Figure S1:**
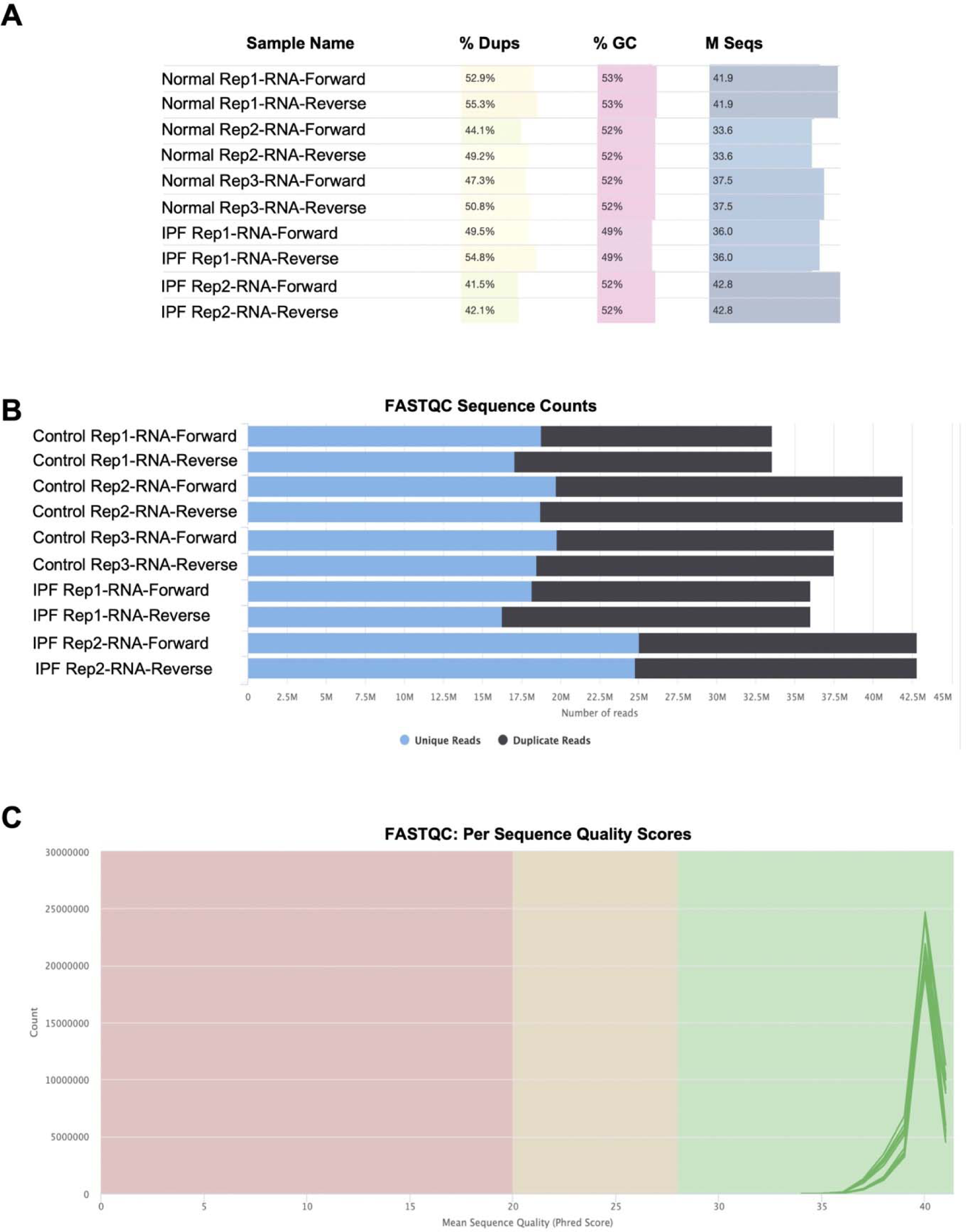
Bulk RNA-seq raw data sequence quality for AT2-control and AT2-IPF samples. **A.** MultiQC analysis of raw RNA FASTQ sequence composition. **B.** MultiQC alignment uniqueness of raw RNA FASTQ reads. **C.** MultiQC quality scores for raw RNA FASTQ sequencing reads.

**Figure S2:**
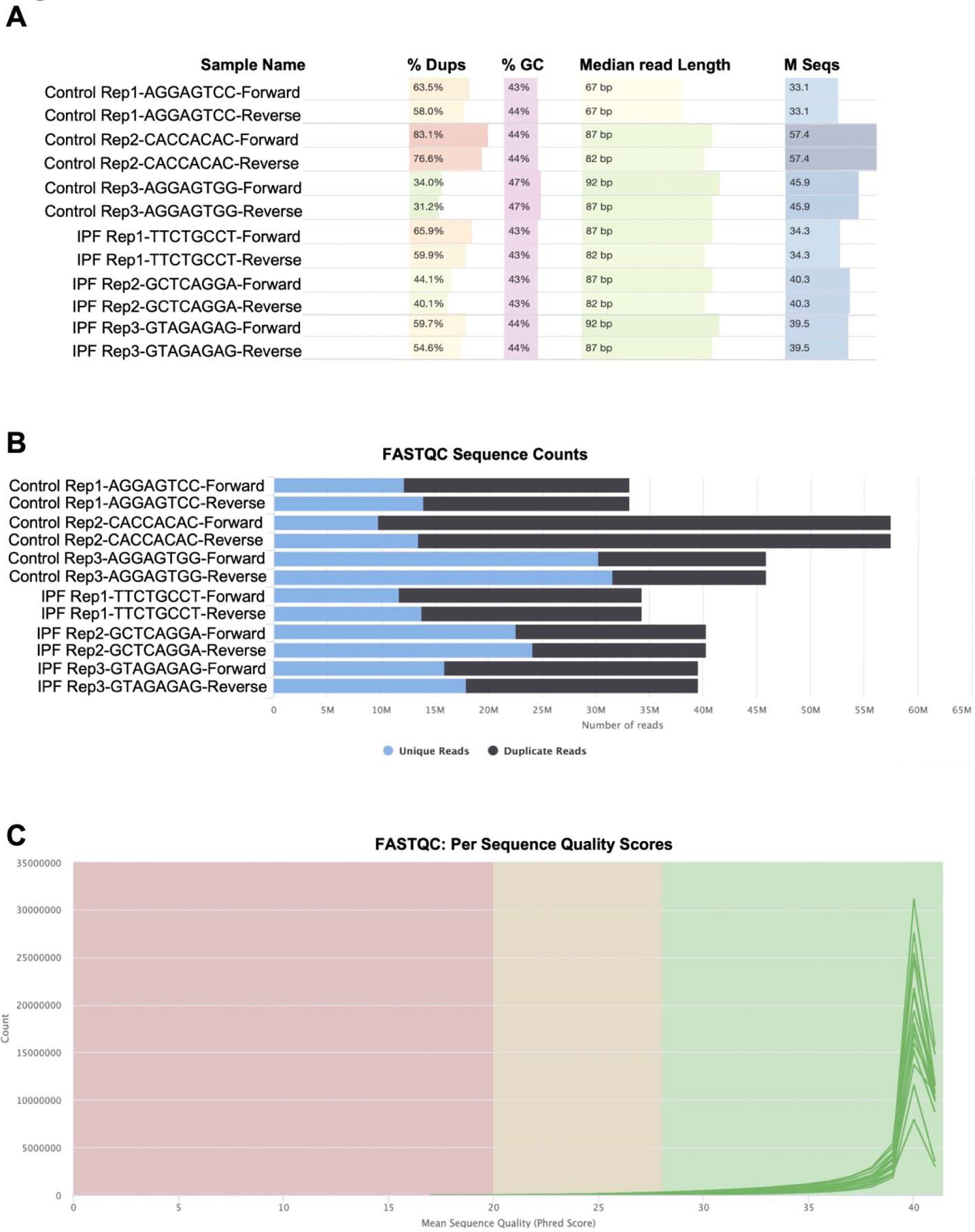
Bulk ATAC-seq raw data sequence quality for AT2-control and AT2-IPF samples. **A.** MultiQC analysis of raw ATAC FASTQ sequence composition. Unique barcodes used for multiplexing are indicated in sample name. **B.** MultiQC alignment uniqueness of raw ATAC FASTQ reads. **C.** MultiQC quality scores for raw ATAC FASTQ sequencing reads.

**Figure S3:**
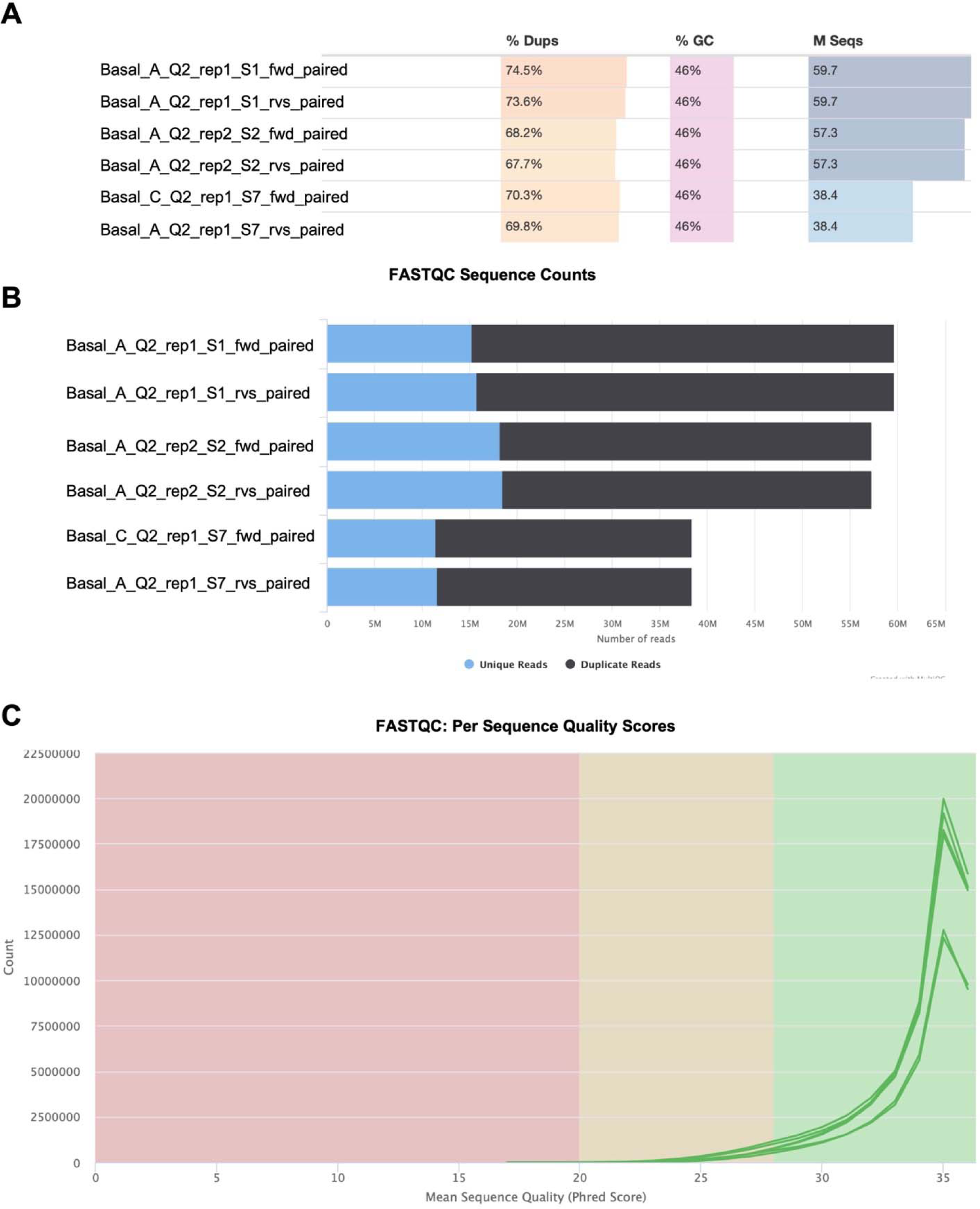
Bulk ATAC-seq raw data sequence quality for basal cells. **A.** MultiQC analysis of raw ATAC FASTQ sequence composition. Unique barcodes used for multiplexing are indicated in sample name. **B.** MultiQC alignment uniqueness of raw ATAC FASTQ reads. **C.** MultiQC quality scores for raw ATAC FASTQ sequencing reads.

**Figure S4:**
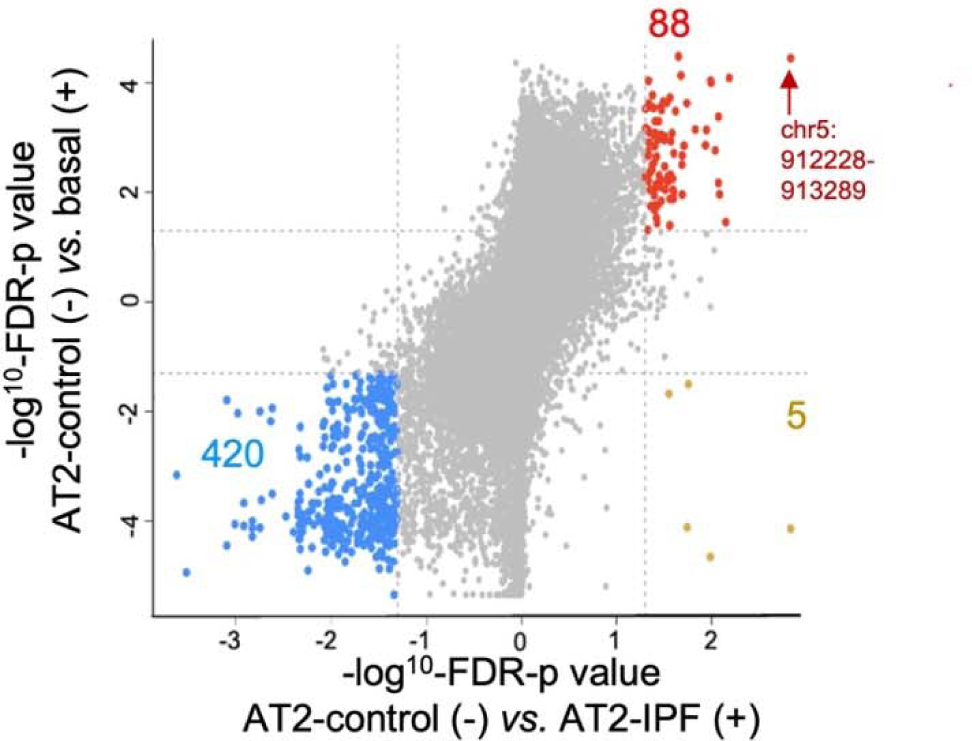
Genome-wide comparison between AT2-IPF, AT2-control and normal basal cells. Starburst plot showing significant peak enrichment based on disease state. X-axis = AT2-IPF (+) vs. AT2-*control* (-). Y-axis = basal (+) vs. AT2-control (-). Red points (88 peaks) = enriched in basal and AT2-IPF relative to AT2-control, light brown (5 peaks) = enriched solely in AT2-IPF, blue (420 peaks) = enriched in AT2-control but not in basal or AT2-IPF, grey = non-significant.

**Figure S5:**
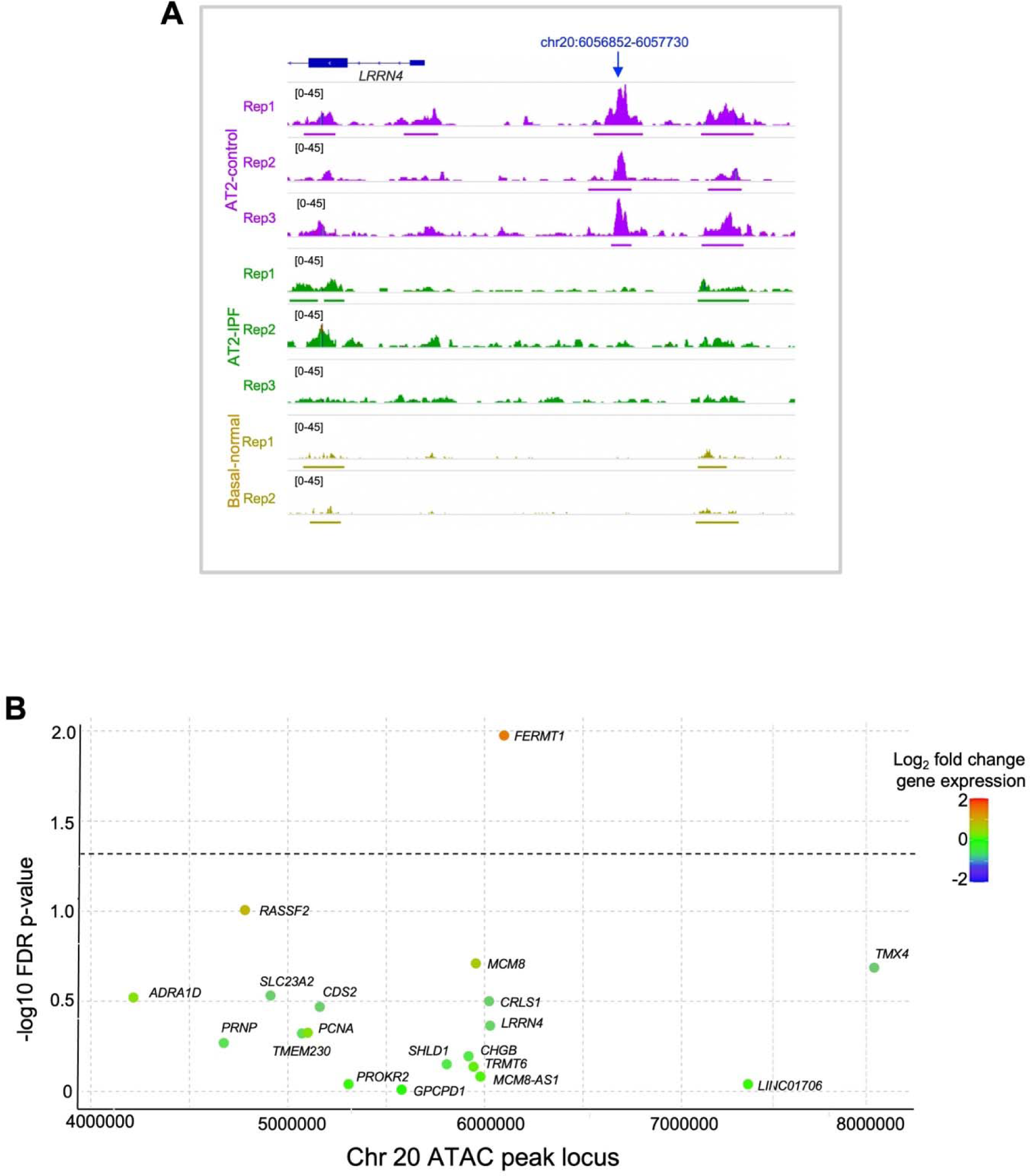
Epigenomic and transcriptomic environment around chr20 peak lost in IPF. **A.** Read count enrichment and called peaks at the chromosome 20 genomic locus where a peak is lost in AT2-IPF, displayed using Integrated Genomics Viewer (IGV). X-axis = Chromosome 20: locus (hg38-based), Y-axis = peak height. AT2-control replicates = purple, AT2-IPF replicates = green, basal replicates = brown. **B.** Local Manhattan plot of the chromosome 20 region 2 Mb to each side of the lost ATAC peak. X-axis = genomic coordinates. Y-axis = -log of p-value for significance in gene expression changes. Dashed line represents significance threshold. Each dot represents a known gene within the examined genomic region. Dot color reflects log2 fold change in gene expression between AT2-IPF and AT2-control. Red = elevated expression in AT2-IPF samples relative to AT2-control, blue = decreased expression in AT2-IPF samples relative to AT2-control. No significantly downregulated gene was observed

**Figure S6:**
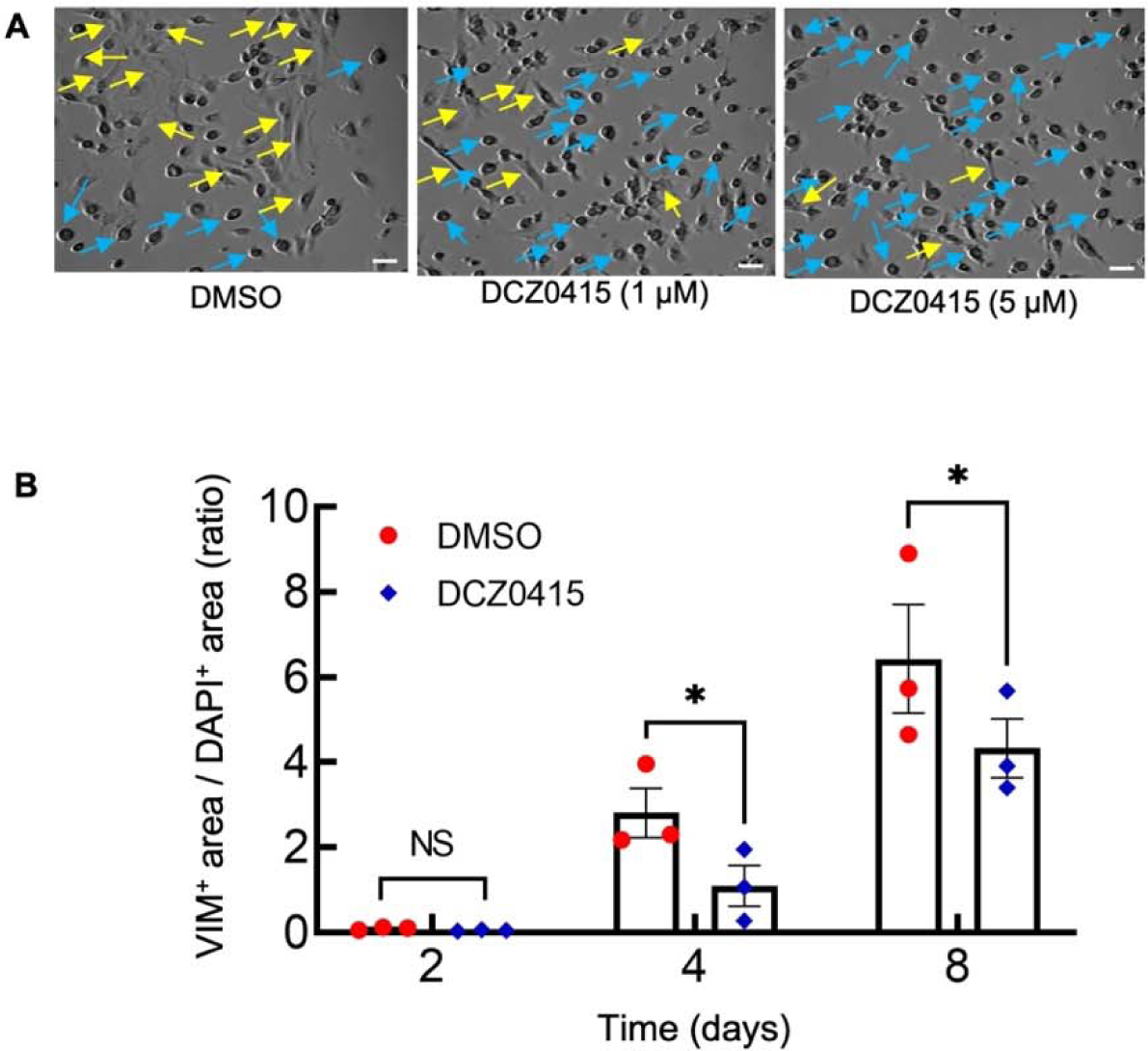
Effect of DCZ0415 on primary human AT2 cells. **A.** Representative phase contrast images of primary human AT2 cells cultured in 2D on plastic treated with vehicle (DMSO) or DCZ0415 for 4 days (light blue arrows indicate cuboidal cells and yellow arrows indicate more elongated fibroblast-like cells) (n = 2). Scale bars: 100 µm. B. Quantitation of area ratio of IF staining of AT2 cells cultured in 2D on glass slides treated with DMSO or DCZ0415 for 2, 4 and 8 days and stained for VIM (n=3) (corresponding to Figure 4E).

## ACKNOWLEDGMENTS

The authors thank the laboratory of Neil Siegel, PhD, for training and advice on ATAC methods. This study was funded by the National Heart, Lung, and Blood Institute (NHLBI) [R35 HL135747 (ZB) and R01HL114959 (BZ)], the Tobacco Related Disease Research Program [T31IP913 (IAO in support of LSP)], the Hastings Center for Pulmonary Research (HCPR) at USC (AB), the Pulmonary Fibrosis Foundation (AB), the Cystic Fibrosis Foundation, CFFT FIRTH21XX0 (ALR, BC, HY, ES and CNM) and the NIH/NCI Norris Comprehensive Cancer Center core grant P30CA014089.

## AUTHOR CONTRIBUTIONS

BZ, CNM, IAO and ZB conceived the work. LSP, ALR, BZ, CNM, ZB and IAO supervised, designed all experiments, and interpreted data. JRA, HJW, YL, YJ, and BZ isolated AT2 cells and performed quality control analysis. LSP, ES, and CNM performed bioinformatic analysis. AB performed TRIP13 validation studies. ALR, BC and HY performed isolation, quality control, and sequencing of primary basal cells. CNM, AB, BZ, IAO, and ZB wrote the manuscript. All authors contributed to the experiments and manuscript finalization.

## INCLUSION AND DIVERSITY

One or more of the authors self-identifies as an underrepresented ethnic minority in science. One or more of the authors self-identifies as a member of the LGBTQIA+ community.

## REFERENCES

1. King TE, Pardo A, Selman M. Idiopathic pulmonary fibrosis. Lancet. 2011;378(9807):1949–61.

2. Luo W, Gu Y, Fu S, Wang J, Zhang J, Wang Y. Emerging opportunities to treat idiopathic pulmonary fibrosis: Design, discovery, and optimizations of small-molecule drugs targeting fibrogenic pathways. Eur J Med Chem. 2023;260:115762.

3. George PM, Patterson CM, Reed AK, Thillai M. Lung transplantation for idiopathic pulmonary fibrosis. Lancet Respir Med. 2019;7(3):271–82.

4. Selvarajah B, Platé M, Chambers RC. Pulmonary fibrosis: Emerging diagnostic and therapeutic strategies. Mol Aspects Med. 2023;94:101227.

5. Pauchet A, Chaussavoine A, Pairon JC, Gabillon C, Didier A, Baldi I, et al. Idiopathic pulmonary fibrosis: What do we know about the role of occupational and environmental Determinants? A systematic literature review and meta-analysis. J Toxicol Environ Health B Crit Rev. 2022;25(7):372–92.

6. Moll M, Peljto AL, Kim JS, Xu H, Debban CL, Chen X, et al. A polygenic risk score for idiopathic pulmonary fibrosis and interstitial lung abnormalities. Am J Respir Crit Care Med. 2023;208(7):791–801.

7. Tirelli C, Pesenti C, Miozzo M, Mondoni M, Fontana L, Centanni S. The genetic and epigenetic footprint in idiopathic pulmonary fibrosis and familial pulmonary fibrosis: A State-of-the-Art Review. Diagnostics (Basel). 2022;12(12).

8. Nogee LM, Dunbar AE, Wert SE, Askin F, Hamvas A, Whitsett JA. A mutation in the surfactant protein C gene associated with familial interstitial lung disease. N Engl J Med. 2001;344(8):573–9.

9. Thomas AQ, Lane K, Phillips J, Prince M, Markin C, Speer M, et al. Heterozygosity for a surfactant protein C gene mutation associated with usual interstitial pneumonitis and cellular nonspecific interstitial pneumonitis in one kindred. Am J Respir Crit Care Med. 2002;165(9):1322–8.

10. van Moorsel CH, van Oosterhout MF, Barlo NP, de Jong PA, van der Vis JJ, Ruven HJ, et al. Surfactant protein C mutations are the basis of a significant portion of adult familial pulmonary fibrosis in a dutch cohort. Am J Respir Crit Care Med. 2010;182(11):1419–25.

11. Wang Y, Kuan PJ, Xing C, Cronkhite JT, Torres F, Rosenblatt RL, et al. Genetic defects in surfactant protein A2 are associated with pulmonary fibrosis and lung cancer. Am J Hum Genet. 2009;84(1):52–9.

12. Campo I, Zorzetto M, Mariani F, Kadija Z, Morbini P, Dore R, et al. A large kindred of pulmonary fibrosis associated with a novel ABCA3 gene variant. Respir Res. 2014;15(1):43.

13. Confalonieri P, Volpe MC, Jacob J, Maiocchi S, Salton F, Ruaro B, et al. Regeneration or repair? The role of alveolar epithelial cells in the pathogenesis of idiopathic pulmonary fibrosis (IPF). Cells. 2022;11(13).

14. Alder JK, Armanios M. Telomere-mediated lung disease. Physiol Rev. 2022;102(4):1703–20.

15. Liu YY, Shi Y, Liu Y, Pan XH, Zhang KX. Telomere shortening activates TGF-β/Smads signaling in lungs and enhances both lipopolysaccharide and bleomycin-induced pulmonary fibrosis. Acta Pharmacol Sin. 2018;39(11):1735–45.

16. Gauldie J, Kolb M, Ask K, Martin G, Bonniaud P, Warburton D. Smad3 signaling involved in pulmonary fibrosis and emphysema. Proc Am Thorac Soc. 2006;3(8):696–702.

17. Ye Z, Hu Y. TGFl1Jβ1: Gentlemanly orchestrator in idiopathic pulmonary fibrosis (Review). Int J Mol Med. 2021;48(1).

18. Bergeron A, Soler P, Kambouchner M, Loiseau P, Milleron B, Valeyre D, et al. Cytokine profiles in idiopathic pulmonary fibrosis suggest an important role for TGF-beta and IL-10. Eur Respir J. 2003;22(1):69–76.

19. Xu Y, Mizuno T, Sridharan A, Du Y, Guo M, Tang J, et al. Single-cell RNA sequencing identifies diverse roles of epithelial cells in idiopathic pulmonary fibrosis. JCI Insight. 2016;1(20):e90558.

20. Adams TS, Schupp JC, Poli S, Ayaub EA, Neumark N, Ahangari F, et al. Single-cell RNA-seq reveals ectopic and aberrant lung-resident cell populations in idiopathic pulmonary fibrosis. Sci Adv. 2020;6(28):eaba1983.

21. Kathiriya JJ, Wang C, Zhou M, Brumwell A, Cassandras M, Le Saux CJ, et al. Human alveolar type 2 epithelium transdifferentiates into metaplastic KRT5. Nat Cell Biol. 2022;24(1):10–23.

22. Carraro G, Stripp BR. Insights gained in the pathology of lung disease through single-cell transcriptomics. J Pathol. 2022;257(4):494–500.

23. Booth LN, Brunet A. The aging epigenome. Mol Cell. 2016;62(5):728–44.

24. Peters A, Nawrot TS, Baccarelli AA. Hallmarks of environmental insults. Cell. 2021;184(6):1455–68.

25. Scruggs AM, Grabauskas G, Huang SK. The Role of KCNMB1 and BK Channels in Myofibroblast Differentiation and Pulmonary Fibrosis. Am J Respir Cell Mol Biol. 2020;62(2):191–203.

26. Sanders YY, Pardo A, Selman M, Nuovo GJ, Tollefsbol TO, Siegal GP, et al. Thy-1 promoter hypermethylation: a novel epigenetic pathogenic mechanism in pulmonary fibrosis. Am J Respir Cell Mol Biol. 2008;39(5):610–8.

27. Korfei M, Mahavadi P, Guenther A. Targeting histone deacetylases in idiopathic pulmonary fibrosis: A Future Therapeutic Option. Cells. 2022;11(10).

28. Zhou B, Stueve TR, Mihalakakos EA, Miao L, Mullen D, Wang Y, et al. Comprehensive epigenomic profiling of human alveolar epithelial differentiation identifies key epigenetic states and transcription factor co-regulatory networks for maintenance of distal lung identity. BMC Genomics. 2021;22(1):906.

29. Marconett CN, Zhou B, Rieger ME, Selamat SA, Dubourd M, Fang X, et al. Integrated transcriptomic and epigenomic analysis of primary human lung epithelial cell differentiation. PLoS Genet. 2013;9(6):e1003513.

30. Liberti DC, Zepp JA, Bartoni CA, Liberti KH, Zhou S, Lu M, et al. Dnmt1 is required for proximal-distal patterning of the lung endoderm and for restraining alveolar type 2 cell fate. Dev Biol. 2019;454(2):108–17.

31. Wang F, Ting C, Riemondy KA, Douglas M, Foster K, Patel N, et al. Regulation of epithelial transitional states in murine and human pulmonary fibrosis. J Clin Invest. 2023;133(22).

32. Love MI, Huber W, Anders S. Moderated estimation of fold change and dispersion for RNA-seq data with DESeq2. Genome Biol. 2014;15(12):550.

33. Donato R, Cannon BR, Sorci G, Riuzzi F, Hsu K, Weber DJ, et al. Functions of S100 proteins. Curr Mol Med. 2013;13(1):24–57.

34. Heukels P, Moor CC, von der Thüsen JH, Wijsenbeek MS, Kool M. Inflammation and immunity in IPF pathogenesis and treatment. Respir Med. 2019;147:79–91.

35. Pardo A, Cabrera S, Maldonado M, Selman M. Role of matrix metalloproteinases in the pathogenesis of idiopathic pulmonary fibrosis. Respir Res. 2016;17:23.

36. Betensley A, Sharif R, Karamichos D. A systematic review of the role of dysfunctional wound healing in the pathogenesis and treatment of idiopathic pulmonary fibrosis. J Clin Med. 2016;6(1).

37. Ponnusamy MP, Seshacharyulu P, Lakshmanan I, Vaz AP, Chugh S, Batra SK. Emerging role of mucins in epithelial to mesenchymal transition. Curr Cancer Drug Targets. 2013;13(9):945–56.

38. Marimuthu S, Rauth S, Ganguly K, Zhang C, Lakshmanan I, Batra SK, et al. Mucins reprogram stemness, metabolism and promote chemoresistance during cancer progression. Cancer Metastasis Rev. 2021;40(2):575–88.

39. Heinz S, Benner C, Spann N, Bertolino E, Lin YC, Laslo P, et al. Simple combinations of lineage-determining transcription factors prime cis-regulatory elements required for macrophage and B cell identities. Mol Cell. 2010;38(4):576–89.

40. Gaspar J. Genrich: Detecting sites of genomic enrichment. Available at https://github.com/jsh58/Genrich. 2018.

41. Ross-Innes CS, Stark R, Teschendorff AE, Holmes KA, Ali HR, Dunning MJ, et al. Differential oestrogen receptor binding is associated with clinical outcome in breast cancer. Nature. 2012;481(7381):389–93.

42. Fraser P, Bickmore W. Nuclear organization of the genome and the potential for gene regulation. Nature. 2007;447(7143):413–7.

43. Smirnova NF, Schamberger AC, Nayakanti S, Hatz R, Behr J, Eickelberg O. Detection and quantification of epithelial progenitor cell populations in human healthy and IPF lungs. Respir Res. 2016;17(1):83.

44. Habermann AC, Gutierrez AJ, Bui LT, Yahn SL, Winters NI, Calvi CL, et al. Single-cell RNA sequencing reveals profibrotic roles of distinct epithelial and mesenchymal lineages in pulmonary fibrosis. Sci Adv. 2020;6(28):eaba1972.

45. Kurita K, Maeda M, Mansour MA, Kokuryo T, Uehara K, Yokoyama Y, et al. TRIP13 is expressed in colorectal cancer and promotes cancer cell invasion. Oncol Lett. 2016;12(6):5240–6.

46. Agarwal S, Afaq F, Bajpai P, Kim HG, Elkholy A, Behring M, et al. DCZ0415, a small-molecule inhibitor targeting TRIP13, inhibits EMT and metastasis via inactivation of the FGFR4/STAT3 axis and the Wnt/β-catenin pathway in colorectal cancer. Mol Oncol. 2022;16(8):1728–45.

47. Lu R, Zhou Q, Ju L, Chen L, Wang F, Shao J. Upregulation of TRIP13 promotes the malignant progression of lung cancer via the EMT pathway. Oncol Rep. 2021;46(2).

48. Willis BC, Liebler JM, Luby-Phelps K, Nicholson AG, Crandall ED, du Bois RM, et al. Induction of epithelial-mesenchymal transition in alveolar epithelial cells by transforming growth factor-beta1: potential role in idiopathic pulmonary fibrosis. Am J Pathol. 2005;166(5):1321–32.

49. Kim KK, Kugler MC, Wolters PJ, Robillard L, Galvez MG, Brumwell AN, et al. Alveolar epithelial cell mesenchymal transition develops in vivo during pulmonary fibrosis and is regulated by the extracellular matrix. Proc Natl Acad Sci U S A. 2006;103(35):13180–5.

50. Marmai C, Sutherland RE, Kim KK, Dolganov GM, Fang X, Kim SS, et al. Alveolar epithelial cells express mesenchymal proteins in patients with idiopathic pulmonary fibrosis. Am J Physiol Lung Cell Mol Physiol. 2011;301(1):L71–8.

51. Goldmann T, Zissel G, Watz H, Drömann D, Reck M, Kugler C, et al. Human alveolar epithelial cells type II are capable of TGFβ-dependent epithelial-mesenchymal-transition and collagen-synthesis. Respir Res. 2018;19(1):138.

52. Sun Y, Jing P, Gan H, Wang X, Zhu X, Fan J, et al. Evaluation of an ex vivo fibrogenesis model using human lung slices prepared from small tissues. Eur J Med Res. 2023;28(1):143.

53. Roach KM, Sutcliffe A, Matthews L, Elliott G, Newby C, Amrani Y, et al. A model of human lung fibrogenesis for the assessment of anti-fibrotic strategies in idiopathic pulmonary fibrosis. Sci Rep. 2018;8(1):342.

54. Alsafadi HN, Staab-Weijnitz CA, Lehmann M, Lindner M, Peschel B, Königshoff M, et al. An ex vivo model to induce early fibrosis-like changes in human precision-cut lung slices. Am J Physiol Lung Cell Mol Physiol. 2017;312(6):L896–L902.

55. Lang NJ, Gote-Schniering J, Porras-Gonzalez D, Yang L, De Sadeleer LJ, Jentzsch RC, et al. Ex vivo tissue perturbations coupled to single-cell RNA-seq reveal multilineage cell circuit dynamics in human lung fibrogenesis. Sci Transl Med. 2023;15(725):eadh0908.

56. Miniowitz-Shemtov S, Eytan E, Kaisari S, Sitry-Shevah D, Hershko A. Mode of interaction of TRIP13 AAA-ATPase with the Mad2-binding protein p31comet and with mitotic checkpoint complexes. Proc Natl Acad Sci U S A. 2015;112(37):11536–40.

57. Yao J, Zhang X, Li J, Zhao D, Gao B, Zhou H, et al. Silencing TRIP13 inhibits cell growth and metastasis of hepatocellular carcinoma by activating of TGF-β1/smad3. Cancer Cell Int. 2018;18:208.

58. Rodríguez A, Epperly M, Filiatrault J, Velázquez M, Yang C, McQueen K, et al. TGFβ pathway is required for viable gestation of Fanconi anemia embryos. PLoS Genet. 2022;18(11):e1010459.

59. Li ZH, Lei L, Fei LR, Huang WJ, Zheng YW, Yang MQ, et al. TRIP13 promotes the proliferation and invasion of lung cancer cells via the Wnt signaling pathway and epithelial-mesenchymal transition. J Mol Histol. 2021;52(1):11–20.

60. Wang Y, Huang J, Li B, Xue H, Tricot G, Hu L, et al. A small-molecule inhibitor targeting TRIP13 suppresses multiple myeloma progression. Cancer Res. 2020;80(3):536–48.

61. Clairmont CS, Sarangi P, Ponnienselvan K, Galli LD, Csete I, Moreau L, et al. TRIP13 regulates DNA repair pathway choice through REV7 conformational change. Nat Cell Biol. 2020;22(1):87–96.

62. Wang K, Sturt-Gillespie B, Hittle JC, Macdonald D, Chan GK, Yen TJ, et al. Thyroid hormone receptor interacting protein 13 (TRIP13) AAA-ATPase is a novel mitotic checkpoint-silencing protein. J Biol Chem. 2014;289(34):23928–37.

63. Pelikan RC, Iwata J, Suzuki A, Chai Y, Hacia JG. Identification of candidate downstream targets of TGFβ signaling during palate development by genome-wide transcript profiling. J Cell Biochem. 2013;114(4):796–807.

64. Jayachandran A, Königshoff M, Yu H, Rupniewska E, Hecker M, Klepetko W, et al. SNAI transcription factors mediate epithelial-mesenchymal transition in lung fibrosis. Thorax. 2009;64(12):1053–61.

65. Yue X, Shan B, Lasky JA. TGF-β: Titan of Lung Fibrogenesis. Curr Enzym Inhib. 2010;6(2).

66. Epstein Shochet G, Brook E, Bardenstein-Wald B, Shitrit D. TGF-β pathway activation by idiopathic pulmonary fibrosis (IPF) fibroblast derived soluble factors is mediated by IL-6 trans-signaling. Respir Res. 2020;21(1):56.

67. Enomoto Y, Katsura H, Fujimura T, Ogata A, Baba S, Yamaoka A, et al. Autocrine TGF-β-positive feedback in profibrotic AT2-lineage cells plays a crucial role in non-inflammatory lung fibrogenesis. Nat Commun. 2023;14(1):4956.

68. Choi J, Park JE, Tsagkogeorga G, Yanagita M, Koo BK, Han N, et al. Inflammatory signals induce AT2 cell-derived damage-associated transient progenitors that mediate alveolar regeneration. Cell Stem Cell. 2020;27(3):366–82.e7.

69. Venkatesan N, Pini L, Ludwig MS. Changes in Smad expression and subcellular localization in bleomycin-induced pulmonary fibrosis. Am J Physiol Lung Cell Mol Physiol. 2004;287(6):L1342–7.

70. Warburton D, Shi W, Xu B. TGF-β-Smad3 signaling in emphysema and pulmonary fibrosis: an epigenetic aberration of normal development? Am J Physiol Lung Cell Mol Physiol. 2013;304(2):L83–5.

71. Zhao J, Shi W, Wang YL, Chen H, Bringas P, Datto MB, et al. Smad3 deficiency attenuates bleomycin-induced pulmonary fibrosis in mice. Am J Physiol Lung Cell Mol Physiol. 2002;282(3):L585–93.

72. Xi Q, He W, Zhang XH, Le HV, Massagué J. Genome-wide impact of the BRG1 SWI/SNF chromatin remodeler on the transforming growth factor beta transcriptional program. J Biol Chem. 2008;283(2):1146–55.

73. Ross S, Cheung E, Petrakis TG, Howell M, Kraus WL, Hill CS. Smads orchestrate specific histone modifications and chromatin remodeling to activate transcription. EMBO J. 2006;25(19):4490–502.

74. Morikawa M, Koinuma D, Miyazono K, Heldin CH. Genome-wide mechanisms of Smad binding. Oncogene. 2013;32(13):1609–15.

75. Kulkarni T, de Andrade J, Zhou Y, Luckhardt T, Thannickal VJ. Alveolar epithelial disintegrity in pulmonary fibrosis. Am J Physiol Lung Cell Mol Physiol. 2016;311(2):L185–91.

76. Eferl R, Hasselblatt P, Rath M, Popper H, Zenz R, Komnenovic V, et al. Development of pulmonary fibrosis through a pathway involving the transcription factor Fra-2/AP-1. Proc Natl Acad Sci U S A. 2008;105(30):10525–30.

77. Wu Q, Zhang KJ, Jiang SM, Fu L, Shi Y, Tan RB, et al. p53: A Key Protein That Regulates Pulmonary Fibrosis. Oxid Med Cell Longev. 2020;2020:6635794.

78. Rana T, Jiang C, Banerjee S, Yi N, Zmijewski JW, Liu G, et al. PAI-1 Regulation of p53 Expression and Senescence in Type II Alveolar Epithelial Cells. Cells. 2023;12(15).

79. Yao C, Guan X, Carraro G, Parimon T, Liu X, Huang G, et al. Senescence of alveolar type 2 cells drives progressive pulmonary fibrosis. Am J Respir Crit Care Med. 2021;203(6):707–17.

80. Picelli S, Björklund AK, Reinius B, Sagasser S, Winberg G, Sandberg R. Tn5 transposase and tagmentation procedures for massively scaled sequencing projects. Genome Res. 2014;24(12):2033–40.

81. Hulsen T, de Vlieg J, Alkema W. BioVenn - a web application for the comparison and visualization of biological lists using area-proportional Venn diagrams. BMC Genomics. 2008;9:488.

82. Thomas PD, Ebert D, Muruganujan A, Mushayahama T, Albou LP, Mi H. PANTHER: Making genome-scale phylogenetics accessible to all. Protein Sci. 2022;31(1):8–22.

83. Dobin A, Davis CA, Schlesinger F, Drenkow J, Zaleski C, Jha S, et al. STAR: ultrafast universal RNA-seq aligner. Bioinformatics. 2013;29(1):15–21.

84. Schneider CA, Rasband WS, Eliceiri KW. NIH Image to ImageJ: 25 years of image analysis. Nat Methods. 2012;9(7):671–5.

85. Bolger AM, Lohse M, Usadel B. Trimmomatic: a flexible trimmer for Illumina sequence data. Bioinformatics. 2014;30(15):2114–20.

86. Fulcher ML, Gabriel S, Burns KA, Yankaskas JR, Randell SH. Well-differentiated human airway epithelial cell cultures. Methods Mol Med. 2005;107:183–206.

87. Buenrostro JD, Giresi PG, Zaba LC, Chang HY, Greenleaf WJ. Transposition of native chromatin for fast and sensitive epigenomic profiling of open chromatin, DNA-binding proteins and nucleosome position. Nat Methods. 2013;10(12):1213–8.

88. Krämer A, Green J, Pollard J, Tugendreich S. Causal analysis approaches in Ingenuity Pathway Analysis. Bioinformatics. 2014;30(4):523–30.

89. Mi H, Muruganujan A, Huang X, Ebert D, Mills C, Guo X, et al. Protocol Update for large-scale genome and gene function analysis with the PANTHER classification system (v.14.0). Nat Protoc. 2019;14(3):703–21.

90. Li H, Durbin R. Fast and accurate short read alignment with Burrows-Wheeler transform. Bioinformatics. 2009;25(14):1754–60.

91. Thorvaldsdóttir H, Robinson JT, Mesirov JP. Integrative Genomics Viewer (IGV): high-performance genomics data visualization and exploration. Brief Bioinform. 2013;14(2):178–92.

92. Gerckens M, Alsafadi HN, Wagner DE, Lindner M, Burgstaller G, Königshoff M. Generation of human 3D lung tissue cultures (3D-LTCs) for disease modeling. J Vis Exp. 2019(144).

93. Horie M, Castaldi A, Sunohara M, Wang H, Ji Y, Liu Y, et al. Integrated single-cell RNA-sequencing analysis of aquaporin 5-expressing mouse lung epithelial cells identifies GPRC5A as a novel validated type I cell surface marker. Cells. 2020;9(11).

94. Khau T, Langenbach SY, Schuliga M, Harris T, Johnstone CN, Anderson RL, et al. Annexin-1 signals mitogen-stimulated breast tumor cell proliferation by activation of the formyl peptide receptors (FPRs) 1 and 2. FASEB J. 2011;25(2):483–96.

